# *In vivo* seamless genetic engineering via CRISPR-triggered single-strand annealing

**DOI:** 10.1101/2022.06.17.496589

**Authors:** Gustavo Aguilar, Milena Bauer, M. Alessandra Vigano, Sophie T. Schnider, Lukas Brügger, Carlos Jiménez-Jiménez, Isabel Guerrero, Markus Affolter

**Author notes:** These authors contributed equally (Gustavo Aguilar, Milena Bauer).

## Abstract

Precise genome engineering is essential for both basic and applied research. CRISPR/Cas accelerated the speed and ease by which defined exogenous sequences are integrated into specific loci. Nevertheless, knock-in generation in multicellular animals remains challenging, partially due to the complexity of insertion screening. Even when achieved, the analysis of protein localization can still be unfeasible in highly packed tissues, where spatial and temporal control of gene labeling would be ideal. Here, we describe SEED/Harvest, an efficient knock-in method based on homology-directed (HDR) and single-strand annealing (SSA) repair pathways. HDR mediates the integration of a switchable cassette. Upon a subsequent CRISPR-triggered repair event, resolved by SSA, the cassette is seamlessly removed. Germline excision of SEED cassettes allows for fast and robust knock-in generation with both fluorescent proteins and short protein tags in tandem. Tissue-specific expression of Cas9 results in somatic cassette excision, conferring spatio-temporal control of protein labelling and the conditional rescue of mutants. Finally, to achieve conditional protein labeling and manipulation of short tag knock-ins, we have developed a toolbox based on rational engineering and functionalization of the ALFA nanobody.

## Introduction

Historically, genetic studies relied on randomly generated mutations. The analysis of such mutants paved the way to the enormous success of genetics in the last hundred years. Yet, as early as the 1980s (Smithies, Gregg, Boggs, Koralewski, & Kucherlapati, 1985), efforts have been devoted to obtain *ad hoc* gene engineering, and thus, rational manipulation of gene products. Today, genetic manipulation is critical for biological studies in all systems. Among other applications, it has permitted the generation of conditional gene knock-outs, gene tagging, and precise base-pair substitutions. Efficient generation of knock-ins has also an immediate application in gene therapy, where it has been successfully employed to correct pathogenic gene variants (DeWitt et al., 2016; So Hyun Park et al., 2019). Hence, the development and implementation of efficient and precise knock-in techniques is, more than ever, urging.

The proposal of CRISPR/Cas as a gene-editing tool (Jinek et al., 2012) revolutionized the generation of knock-ins, both in terms of ease and precision. Briefly, Cas9 enzyme, loaded with specific guideRNAs, is used to generate double strand breaks (DSB) in a target locus, triggering the DNA’s repair machinery. An exogenous DNA containing the intended insertion is provided as a repair template during the process (Ceccaldi, Rondinelli, & D’Andrea, 2016). With some exceptions, Homology-directed repair (HDR) is the preferred pathway used to generate precise knock-ins (Bollen, Post, Koo, & Snippert, 2018). HDR rates are often low *in vivo* (Johnson & Jasin, 2000; Mao, Bozzella, Seluanov, & Gorbunova, 2008), and thus, an efficient way for screening correct insertions is required.

To facilitate the screening, two-step knock-in approaches have grown increasingly popular, especially when editing multicellular organisms. In these cases, a selectable marker is inserted along with the desired DNA element. After screening, the marker is removed by recombination via the Cre/LoxP or Flipase/FRT systems (Bollen et al., 2018), This approaches result in scars of variable sizes that, although sometimes innocuous, are generally incompatible with precise protein tagging. As an alternative, piggyBac (PBac) techniques allow seamless marker removal (Yusa, 2013). This approach depends on the presence of a natural TTAA site around the insertion point or *de novo* engineering of such a site. PBac also requires targeted expression of a transposase and entails the risk of reintegration in other genomic locations (Ye et al., 2014).

Recently, approaches based on Microhomology-dependent End-Joining (MMEJ) permitted seamless removal of a marker from an engineered locus in iPSC cells (Kim et al., 2018; Roberts et al., 2019). In these approaches, the marker is flanked by guide RNA (gRNA) targets at both sides, cleavable by CRISPR/Cas. To favor MMEJ, small repeats (up to 50bp) are inserted at both sides of the gRNA targets. This strategy has been employed in cultured cells to create point mutations or insert fluorescent tags, where it has reached an efficiency of up to 50% (Roberts et al., 2019). The solely need of CRISPR/Cas makes this approach ideal for gene editing in multicellular organisms.

The insertion of a tag at the endogenous locus permits both the visualization and the manipulation of the targeted gene product. Nonetheless, visualization of the protein of interest may remain limited when studying highly-packed tissues, like the brain. To overcome these limitations, genetic engineering that permits endogenous tagging in a distinct subset of cells has been proposed (Alexandre, Baena-Lopez, & Vincent, 2014; Baena-Lopez, Alexandre, Mitchell, Pasakarnis, & Vincent, 2013; Koles, Yeh, & Rodal, 2015). Such approaches often require up to four integration/excision events (Baena-Lopez et al., 2013), making its application in vertebrates costly and time consuming. In addition, these approaches require the implementation of an increasingly complex genetic toolbox, lacking in most species.

Besides genetic approaches, protein-based methods, such as chromobodies, have also been proposed to label proteins in specific cells. Chromobodies are genetically-encoded fusions of a fluorescent protein and a protein binder. Upon chromobody expression, the protein binder selectively recognizes and labels its target. Numerous protein binders, including nanobodies, single-chain antibodies (scFvs) and monobodies have been employed for protein visualization (Gross et al., 2013; Helma et al., 2012; Kamiyama et al., 2015; Panza, Maier, Schmees, Rothbauer, & Soellner, 2015). Often, chromobodies directly detect an epitope on the endogenous protein, which entails the isolation of different protein binders for each new target. Alternatively, chromobodies can target a protein tag. This strategy allows for the generation of universal chromobodies that can be used to detect multiple targets. However, this approach requires endogenous protein tagging, placing again the limiting step in the gene editing.

Here, we propose a two-step approach to rapidly and robustly generate knock-ins in Drosophila. The technique, named SEED (from “**S**carless **E**diting by **E**lement **D**eletion”)/Harvest, relies only on CRISPR/Cas for both steps. SEED exploits HDR and Single stranded annealing (SSA) repair pathways to insert and seamlessly remove a screening marker, respectively. SEED/Harvest provides a fast, robust and affordable alternative for current tagging methods. The high efficiency of SSA in somatic cells also permits conditional gene labeling, mediated by the tissue-specific expression of Cas9. This is the first example of lineage-restricted endogenous tagging mediated by CRISPR/Cas. In order to visualize knock-ins with short peptide tags, we have also developed a novel nanobody-based toolbox based on engineering and functionalization of the anti-ALFA nanobody (ALFANb) (Götzke et al., 2019). Together, these reagents permit the tissue-specific visualization and manipulation of endogenous proteins via both genetic- and protein-based methods.

## Results

### SEED/Harvest: Generation of scarless genome insertions via a combination of HDR and SSA repair pathways

We envisioned a scarless knock-in strategy that makes use of the SSA pathway in Drosophila. SSA is a highly conserved repair pathway, strongly preferred when repeated sequences flank the DSB (Bhargava, Onyango, & Stark, 2016; Preston, Engels, & Flores, 2002). In our approach, a cassette is incorporated in the gene of interest via CRISPR-triggered HDR (Figure 1A). The cassette consists of a selectable marker (3×P3-dsRED, (Berghammer, Klingler, & A. Wimmer, 1999)), flanked by the target sequences of two gRNAs with no cutting sites anywhere else in the genome (here named gRNAs #1 and #2) (Garcia-Marques et al., 2019). Flanking these sites, the exogenous sequence to be inserted is split in a 5’ and a 3’ parts, sharing a repeated sequence of 100bp to 400bp. The SEED cassette is then flanked by the 5’ and 3’ homology arms required for HDR-mediated genome insertion. In order to trigger plasmid linearization *in vivo*, the target sequences of the gRNA used to generate the genomic DSB are also added flanking the whole construct. After insertion of the SEED cassette and screening, the screening marker is seamlessly removed by a subsequent CRISPR-triggered repair event, resolved by SSA (Figure 1B). In this step, the repeats anneal and the region in-between is removed, resulting in the scarless removal of the 3×P3-dsRED marker and the desired gene editing (Figure 1C). Our strategy was inspired by the recently developed CRISPR-triggered cell-labeling (Garcia-Marques et al., 2019) and has a pilot precedent in cell culture (Li et al., 2018).

**Figure 1.**
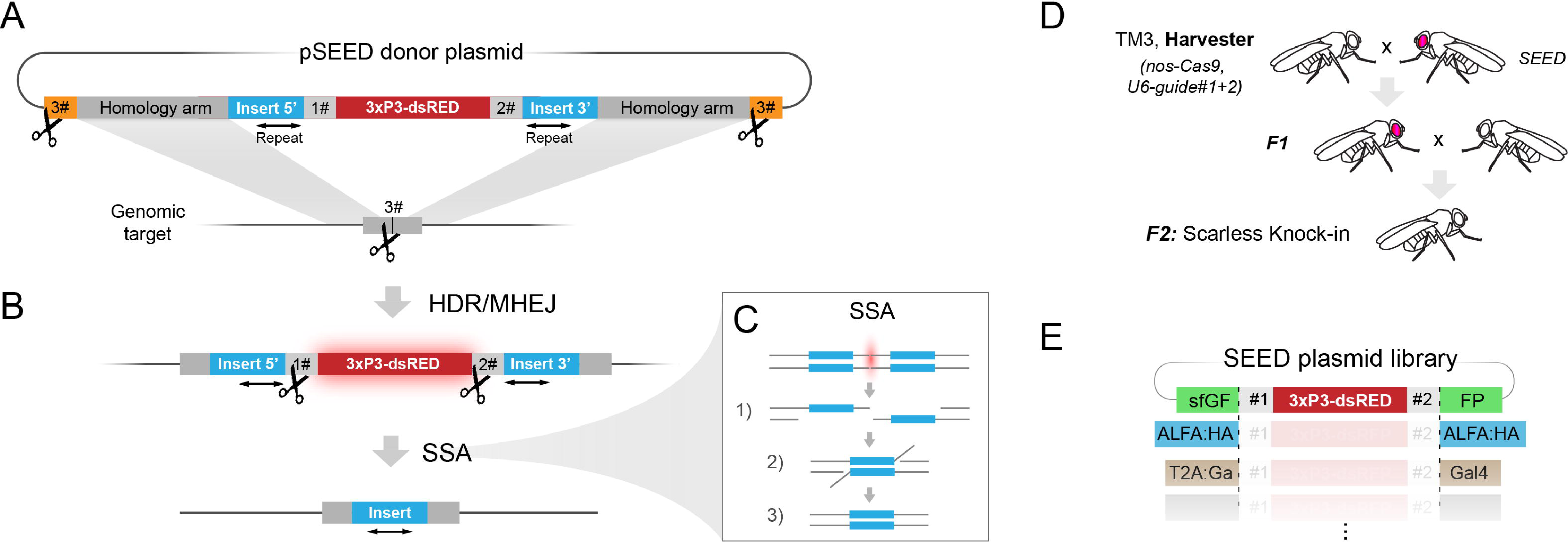
The SEED/Harvest strategy. **A.** Outline of the knock-in strategy. pSEED donor plasmids are composed of the following elements: 1) the selectable marker 3×P3-dsRED (red box), 2) targets for two gRNAs with no cutting sites in the fly genome (in grey), here denominated 1# and 2# (following the nomenclature of Garcia-Marques et al. 2019), 3) the intended sequence to be inserted in the target locus (in blue), this sequence will be split in two parts (“3’ insert” and “5’ insert”), with a common repeat of 100-400 bp, 4) the homology arms to trigger HDR upon DSB formation, and 5) the target sequences of the gRNA that will also cut the genome (in orange), flanking the homology arms (here arbitrarily denominated 3#). Upon providing Cas9 and the gRNA 3#, a DSB will be generated in the locus of interest. The DSB will be then repaired by HDR using the highly recombinogenic linearized donor as a template. **B.** Harvesting step. Upon insertion and selection via the 3xP3-dsRED marker, gRNAs 1# and 2# and Cas9 are provided, resulting in the excision of the dsRED marker. Given the presence of the repeats flanking the DSB, SSA will be the preferred repair pathway, leading to the seamless reconstitution of the full-length desired insert. **C.** Schematic representation of the highly conserved steps of SSA repair; 1) 5’-specific strand degradation, 2) Single strand annealing, 3) Resection of overhangs and ligation. **D.** Example of harvesting crossing scheme. Harvester flies, bearing *nos*-Cas9 *U6*-gRNAs #1+2 inserted in a balancer chromosome are bred with flies containing the desired SEED cassette. In the germline of the F1, the cassette is excised. Outcross of these animals lead to F2 in which the cassette is deleted. (For full information, see Supplementary Figure 1). **E.** Schematic representation of the ready-to-use SEED plasmid library. 3xP3-dsRED is flanked by the target sequences of gRNAs 1# and 2#. These in-turn are flanked by the intended insertion, split in 5’ and 30 regions, and containing a repeat. Here shown sfGFP, ALFA:HA and T2A:Gal4. Complete list in Supplementary Figure 1F.

To facilitate cassette removal and selection, we generated flies containing a balancer chromosome bearing both ubiquitously expressed gRNAs #1 and #2 and a germ line-driven Cas9 (*nos-Cas9, U6-gRNA#1+2*), hereafter named “Harvester” (Supplementary Figure 1A). Harvester stocks permit the generation and selection of knock-ins in one and a half months (Figure 1D, Supp Figure 1E).

In order to quantitatively test the efficiency of the SSA-mediated rearrangement of SEED cassettes *in vivo*, we generated a construct containing a sfGFP-SEED cassette downstream of UAS sequences (*UAS-sfGFP-SEED*) and inserted it in the *attP*86Fb landing site via integrase-mediated insertion. *UAS-sfGFP-SEED* flies displayed easily recognizable dsRED signal in the mid-gut, brain and eyes in both larvae and adults (Supplementary Figure 1B, (Berghammer et al., 1999)). When un-cleaved, expression of *UAS-sfGFP-SEED* results in a non-fluorescent truncated sfGFP protein, due to a stop codon immediately before the 3xP3-FP marker. Only in those cases in which the targeted locus undergoes SSA, the full sfGFP coding sequence is reconstituted. We analyzed the progeny of *Harvester*/*UAS-sfGFP-SEED* flies crossed with the ubiquitous driver *Act5C*-Gal4 (Supplementary Figure 1C, D). Of those flies that inherited the *UAS-sfGFP-SEED* insertion, most of the progeny expressed GFP throughout the body (87.8%). Among the GFP-negative flies, half of them still kept the 3xP3-dsRED marker, permitting their sorting. Thus, of the flies that lost the marker 94.8% underwent seamless rearrangement, the other 5.2% probably being repaired by other pathways. This result is consistent with other reports of SSA frequency in Drosophila (Dewey, Korda Holsclaw, Saghaey, Wittmer, & Sekelsky, 2023; Garcia-Marques et al., 2019) In addition, we tested the efficiency of the ALFA:HA-SEED harvesting. Despite the shorter repeat (100bp), in 67% of the cases, SSA happened seamlessly when 3xP3-dsRED was removed (Supplementary Figure 1D).

We then generated a customized library of SEED cassettes (Figure 1E, Supplementary Figure 1F) ready to be used to insert either popular fluorescent proteins (sfGFP, mCherry, EYFP:HA), recently generated fluorescent proteins with improved brightness (mScarlet2 (Bindels et al., 2017) and mGreenLantern (mGL (Campbell Benjamin et al., 2020)) or short protein tags in tandem (ALFA-tag (Götzke et al., 2019), HA-tag (Wilson et al., 1984), MoonTag (Boersma et al., 2019) and OLLAS-tag (S. H. Park et al., 2008)). Since dsRED and mCherry share long stretches of sequence that might interfere during SSA repair, the 3xP3-dsRED of pSEED-mCherry and pSEED-mScarlet plasmids was substituted for 3xP3-GFP. For all the chosen proteins and short tags, highly selective antibodies, as well as high-affinity protein binders that enable *in vivo* manipulation are available (Aguilar, Vigano, Affolter, & Matsuda, 2019). In addition to tagging reagents, we generated a pSEED-T2A-GAL4 plasmid that allows rapid generation of Gal4 driver lines, and an empty SEED vector, permitting the generation of *ad hoc* SEED cassettes. All plasmids are designed to facilitate one day assembly via Golden Gate cloning (Engler, Kandzia, & Marillonnet, 2008) (Supplementary Figure 2A and Material and methods).

### Precise gene tagging using SEED/Harvest

As a proof of principle, we attempted the tagging of the *patched* (*ptc*) and the *spaghetti squash* (*sqh*) genes with the pSEED-ALFA:HA donor plasmid. In both cases, gRNAs and Cas9 were provided by integrated transgenes, as this approach has been shown to result in enhanced integration efficiency (Port, Chen, Lee, & Bullock, 2014a). Long homology arms (∼1.4kb) were used flanking the cassettes (see Figure 2A for an overview of the strategy). High rates of insertion in the germ cells (Seeding) were observed, with 58% (*ptc*) and 50% (*sqh*) of the F0 parents giving rise to progeny positive for the 3xP3 marker (Figure 2G). Seamless SEED cassette removal (Harvesting) occurred in similar rates as for *UAS-ALFA:HA-SEED* (Figure 2H). Imaginal wing disc immunostaining of Ptc:ALFA:HA permitted simultaneous visualization of both tags using anti-HA monoclonal antibody and fluorescently conjugated anti-ALFA nanobody (Figure 2B and B’). Ptc was detected along the antero-posterior (AP) boundary, in its characteristic anterior stripe (Capdevila, Estrada, Sanchez-Herrero, & Guerrero, 1994). Sqh:ALFA:HA was detected in all cells, decorating the cellular cortex (Figure C).

**Figure 2.**
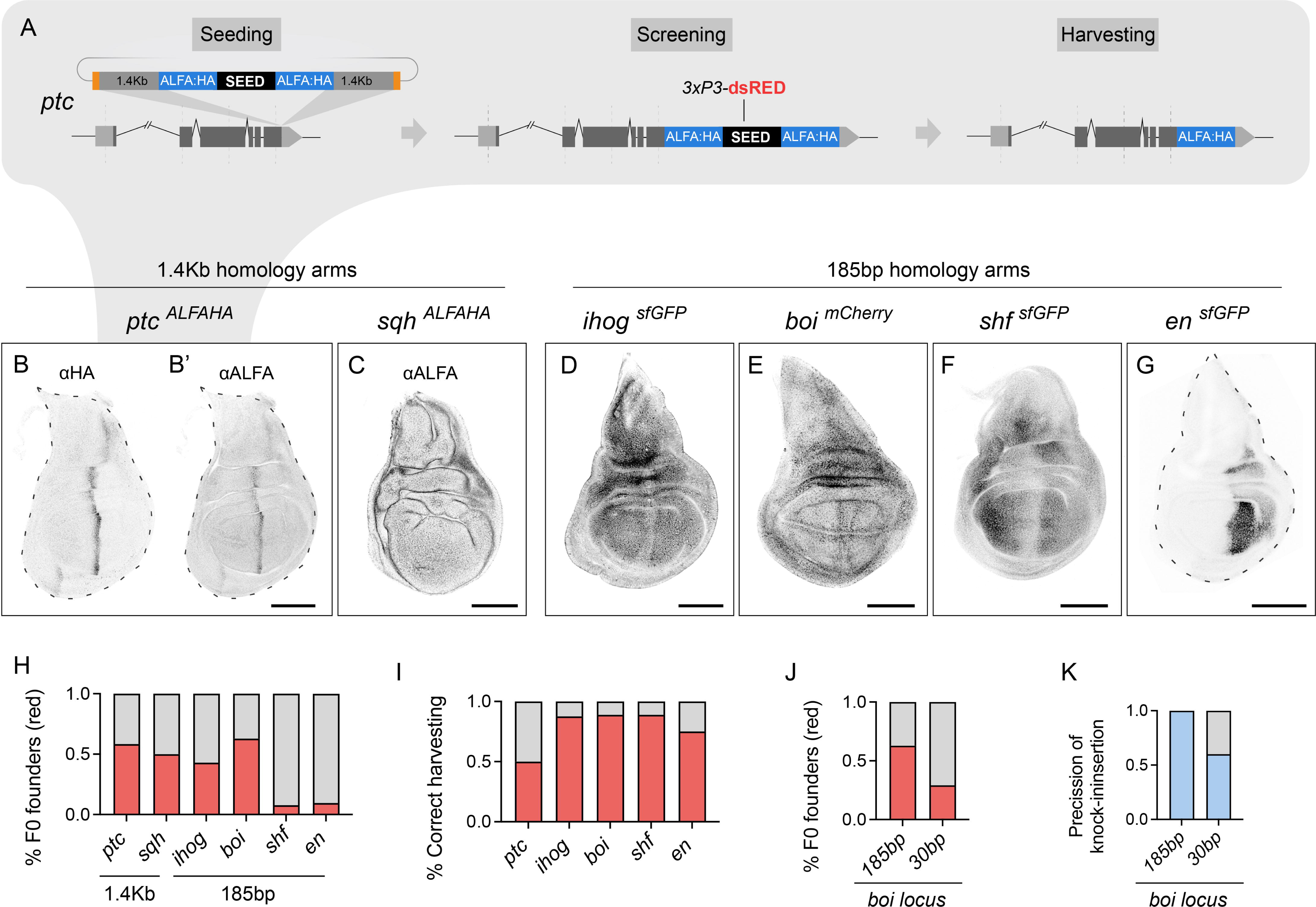
Precise knock-in generation using SEED/Harvest. **A.** Details of the *ptc* knock-in strategy. 1) Seeding. Two gRNAs targeting the vicinity of the STOP codon were utilized to trigger insertion of the SEED-ALFA:HA cassette via CRISPR-triggered HDR. 2) Screening. 3xP3-dsRED animals permitted the isolation of knock-ins, further validated by PCR and Sanger sequencing. 3) Harvesting. Harvester flies (See Figure 1 and Supplementary Figure 2) were used to remove the SEED cassette in a SSA-mediated repair events, which resulted in seamless ALFA:HA insertion. **B.** Ptc:ALFA:HA wing imaginal disc co-stained with an anti-HA antibody and anti-ALFA nanobody (**B.**), (confocal images) The anterior stripe pattern mimicked previously reported expression (Capdevila et al., 1994). **C.** Anti-ALFA nanobody staining of Sqh:ALFA:HA wing disc. Sqh is ubiquitously expressed, with cortical sub-cellular distribution, single confocal plane. **D.** Patterns of Ihog:sfGFP, **E.** Boi;mCherry, **F.** sfGFP:Shf and **G.** sfGFP:En localization in the wing disc after SEED harvesting, In all cases, a single confocal plane of the endogenous fluorescence is shown. Ihog:sfGFP presents its characteristic downregulation along the A-P boundary. Boi:mCherry is being localized in a characteristic cross pattern, sfGFP:Shf is downregulated in the central part in response to Hh, sfGFP:En localizes in the nuclei of the posterior compartment and first cell rows of the anterior wing pouch. **H.** Rates of fertile F0 that gave rise to dsRED-positive progeny (red): *ptc*: 58% (n=60), *sqh*: 50% (n=8), *ihog*: 43% (n=28), *boi*: 63% (n=35), *shf*: 8% (n=24), *en*: 11% (n=76). **I.** Rates of SSA seamless rearrangement of the SEED cassettes (red) among 3xP3-dsRED negative individuals. **J.** Comparison of knock-in efficiency (red) using homology arms of 185bp and 30bp in the *boi* locus. 185bp: 63% (n=25), 30bp 41% (n=39). **K.** Comparison of seamless insertion (integration without indels or mismatches) (light blue) using homology arms of 185bp and 30bp in the *boi* locus. 185bp: 100% (n=10), 30bp 60% (n=10). Scale bars: 100μm.

Often, one of the main limiting steps in the generation of knock-ins is the construction of donor plasmids. Amplification and cloning of long homology arms can complicate or even hamper transgene generation. To increase cloning speed and efficiency, we adopted the use of short homology arms (<200bp) (Kanca et al., 2019). This strategy permits the cheap synthesis of vectors including homology arms (around 80€ by the time this manuscript is written), into which the SEED cassette can be added in a one-day cloning (Supplementary Figure 2A, see material and methods). Using this strategy, we tagged the genes encoding for the Hedgehog (Hh) pathway components: *interference hedgehog* (*ihog*), *brother of ihog* (*boi*) and *shifted* (*shf*), and the transcription factor *engrailed* (*en*) with sfGFP or mCherry (Supplementary Figure 2 E-H). Instead of using genetically provided gRNAs, we co-injected gRNA expressing plasmids together with the donor templates. In two of the cases (*ihog* and *boi*), this strategy resulted in a similar integration efficiency as that observed in the *ptc* locus employing long homology arms and genetically provided gRNAs. (Figure 2H). Knock-ins in *en* and *shf* locus occurred at lower rates (Figure 2H). For all the generated knock-ins, the corresponding expression patterns, revealed by immunostaining or fluorescent detection in wing imaginal discs, mimicked that previously described in the literature (Figure 2D-G) (Bilioni et al., 2013; Brower, 1986; Glise et al., 2005; Yan et al., 2010) and all resulted in functional variants, presenting no obvious mutant phenotypes (data not shown).

Recent studies have proposed the integration of transgenes employing much shorter homology arms, exploiting either the Micro-Homology End Joining (MHEJ) (Nakade et al., 2014; Sakuma, Nakade, Sakane, Suzuki, & Yamamoto, 2016) or more generally homologous-mediated end joining (HMEJ) (Yao et al., 2018) pathways. MHEJ can support transgene integration with sequences as short as 5bp. Since the use of such homology arms could further simplify the generation of the donor, we generated a pSEED-mCherry donor vector bearing 30bp homology arms for the *boi* locus. Thus, donor plasmids could be easily generated in a one-day cloning (Supplementary figure 2B), skipping the synthesis step. Insertion points and gRNAs were identical to those employed in Figure 2D, and can thus be directly compared. The percentage of founder animals among fertile F0 projeny was 41% (Figure 2I), which only represented a slight decrease with respect to the 63% using donors with 185bp homology arms. However, the use of such short homology arms resulted in a decrease of the precision with which the transgenes are integrated; while 185bp resulted in 100% accuracy, presenting no indels or mutations, using 30bp resulted in 60% of the founders containing accurate insertions (Figure 2J). Together, our data suggests that MHEJ-dependent knock-ins represent a promising alternative to HDR in Drosophila.

### Generation of point mutations via SEED/Harvest

In many occasions, especially in gene therapy, the intended genetic manipulation involves not the addition of a tag, but the mutation of one or several base-pairs. With slight modifications, SEED/Harvest can achieve the generation of point mutants. When not introducing a tag, the repeats used for SSA-mediated harvesting have to be added during the cloning, matching the endogenous sequence. This approach has been proven efficient in cell culture (Li et al., 2018), but its application *in vivo* has remained unexplored. As a proof of concept, we have recently used SEED/Harvest to generate a point mutant in the *scalloped* (*sd*) gene, member of the TEAD transcription factor family (Mesrouze et al., 2022) (Supplementary figure 3A). In this case, the region to be mutated was duplicated at both sides of the SEED cassette (Supplementary Figure 3B). These repeats of approximately 100 bp contained the point mutation and silent mutations to avoid donor cutting by the gRNAs. The HDR mediated insertion resulted in a high number of dsRED positive animals (52%, n=33). Furthermore, we could harvest the SEED to generate the desired edited mutant stock (Supplementary Figure 3C).

### CRISPR-triggered lineage-specific endogenous labeling

One of the main limitations of endogenous labeling when studying protein localization is the widespread distribution of many proteins. This turns out to be particularly critical in tissues with an intricated three-dimensional structure, such as the brain, where linage-specific endogenous labeling would be most useful. In order to obtain tissue-specific protein labeling, the most common strategy is the overexpression of a tagged cDNA. Overexpression approaches result in high amounts of protein being synthesized, with the risk of altering protein function and localization. Prior to harvesting, all SEED cassettes encoding fluorescent proteins contain a stop codon after the first repeat, producing a truncated non-fluorescent protein. We hypothesized that these cassettes could be used as switches in somatic cells, responsive to the presence of Cas9 and the relevant gRNAs. We attempted to switch the *ihog*^sfGFP-SEED^ allele in wing disc cells by tissue-specific expression of Cas9 (Figure 3A). gRNAs were provided ubiquitously by a *U6-*gRNA#1+2 transgene. The *UAS*-u^M^Cas9, a transgene with tempered Cas9 expression, was used in all experiments to avoid side effects derived from high Cas9 levels (Port et al., 2020). In contrast to control discs, in which no fluorescence was detected (Figure 3B), expression of Cas9 using the *ap*-Gal4 driver resulted in the restricted labeling of Ihog within the dorsal compartment (Figure 3B’). While most of the compartment is positive for Ihog:sfGFP, some small clones of cells remain unlabeled, suggesting that the SEED cassette is not repaired via SSA or that the cassette remains uncut in these cells. Similarly, expression of Cas9 in the wing pouch using the *nub*-Gal4 driver also resulted in restricted labeling (Fig 3B’’). In these experiments, Ihog:sfGFP positive clones were also often detected outside the driver domain (Figure 3C). It has been reported that the off-target effect of Gal4-driven Cas9 can be attenuated by restricting gRNA expression to the target cells (Port & Bullock, 2016). Compared to the *U6* promoter, *UAS*-driven gRNAs #1 and #2 resulted in drastic reduction of off-target labeling when expressed via *ap-*Gal4 (compare Figure 3C and C’, quantified in D). In order to confirm that somatic switching is extendable to other loci, we used the same approach to trigger sfGFP:En labelling in a *en^sfGFP-SEED^*background. As predicted, no fluorescence was detected in the un-switched allele (Figure 3E). Expression of *UAS*-u^M^Cas9 and *UAS*-gRNAs#1+2 under the *ap*-Gal4 driver resulted in robust En labeling in the dorsal compartment (Figure 3E’). Within the targeted cells, no signal was detected outside the *en* expression domain (Compare to germ-line harvested, *en*^sfGFP^ (Figure 3F)), which confirms that somatic labeling via SEED does not alter the endogenous regulation of gene expression.

**Figure 3.**
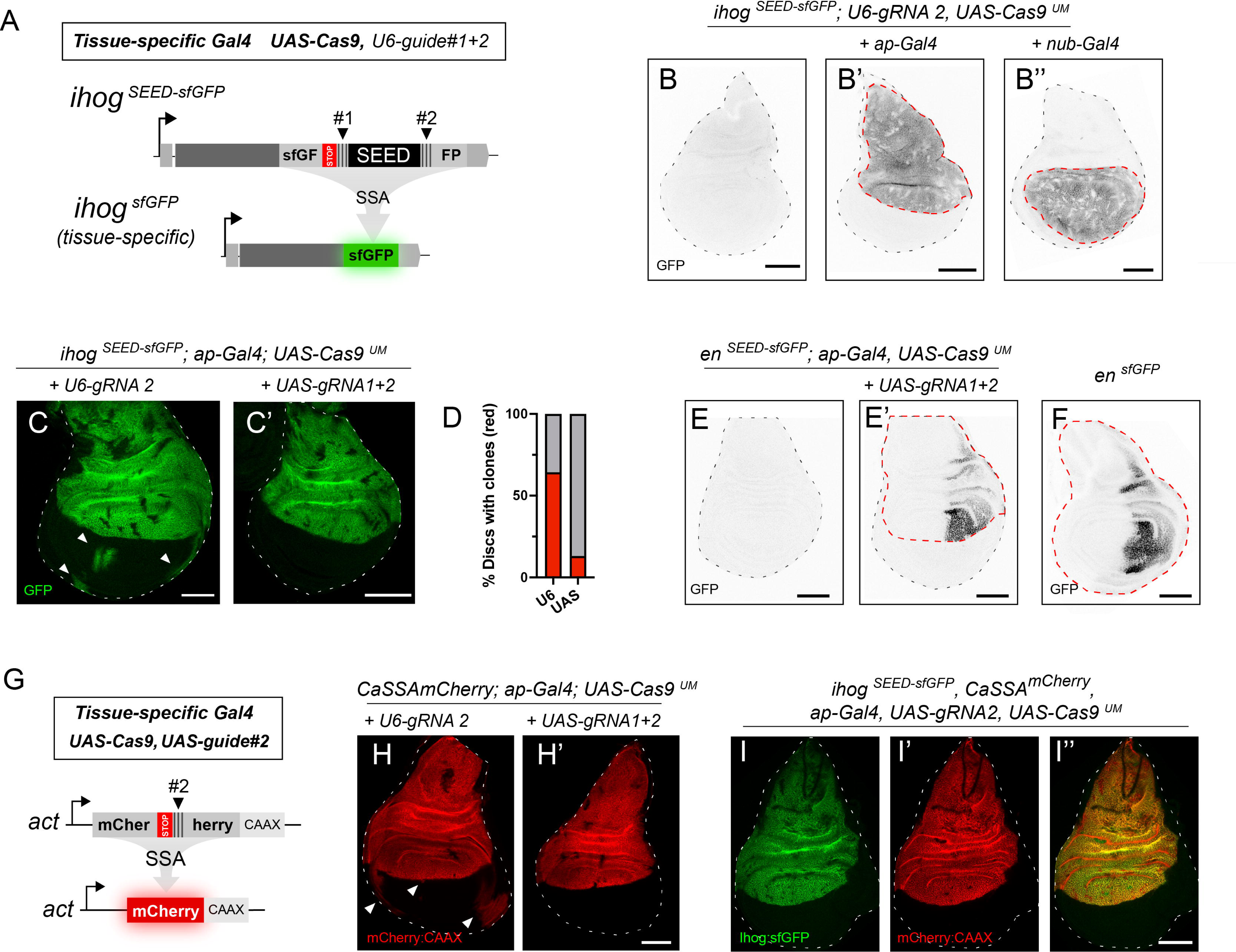
CRISPR-triggered somatic labeling. **A.** Outline of the somatic labeling concept. Flies bearing a sfGFP SEED cassette are crossed with strains that contain Cas9 expressed under a tissue-specific promoter and ubiquitously expressed gRNAs. Before cleavage, proteins are labelled with a truncated version of sfGFP. After DSBs are induced, the sfGFP open reading frame is reconstituted, giving rise to a fluorescent sfGFP-tagged protein. **B**. GFP fluorescence of an *ihog*^SEED-sfGFP^, *U6-gRNA#1+2* wing disc without driver (**B**), driven by *ap*-Gal4 (**B’**) or by *nub*-Gal4 (**B’’**). Most cells within the domain express Ihog:GFP, but Ihog negative clones can be observed. **C.** Example of wing disc displaying clones of Ihog:sfGFP outside the *ap* expression domain when gRNAs are provided using a *U6* promoter. Arrowheads indicate the ectopic clones. **C’.** Tissue-specific harvesting using *UAS*-driven gRNAs. **D.** Quantification of the experiments shown in C and C’. U6-driven n=12, UAS-driven n=25. **E.** *en*^SEED-sfGFP^ control wing disc. **E.** Dorsal harvesting of *en*^SEED-sfGFP^ in the wing disc, using *ap*-Gal4. Dotted red line indicates *ap* expression domain. **F.** Wing disc of an *en*^sfGFP^, generated by germ-line harvesting of *en*^SEED-sfGFP^ using Harvester flies. **G.** Scheme of CaSSA reporters. Act5C promoter drives the expression of a mCherry:CAAX cassette. mCherry is split in a 5’ and 3’ regions, sharing an internal repeat. In between, gRNA#2 target sequences permit the cleavage of the reporter, resulting in SSA and the reconstitution of mCherry. In our approach, Cas9 and gRNAs are provided using a tissue-specific Gal4. **H.** Wing disc in which CaSSAmCherry reporter has been activated in the *ap* domain. Notice the numerous clones outside the dorsal compartment (arrows) when using *U6*-driven gRNAs. **H’.** Activation of CaSSAmCherry in the *ap* domain using *UAS*-driven gRNAs. **I.** Wing disc showing double labeling of endogenous Ihog:GFP (**I**) and cell membranes (**I’**), using SEED and CaSSA respectively, within the *ap* domain. **I’’**. Merge of I and I’. Scale bars: 100μm.

One advantage of the somatic gene labeling via SEED/Harvest is its compatibility with other CRISPR-triggered applications. In recent years, tissue-specific expression of Cas9 has been proposed to achieve cell labeling, tissue-specific mutagenesis or the generation of chromosomal crossovers, among other uses (Allen et al., 2021; Garcia-Marques et al., 2019; Koreman et al., 2021; Meltzer et al., 2019; Port & Bullock, 2016; Port et al., 2020). To favor compatibility, SEED cassettes are designed to be triggered by the same gRNAs (#2) used by that CaSSA reporters. Originally developed for linage labelling in the brain, CaSSA membrane reporters are also based on the reconstitution of a fluorescent protein via CRISPR-triggered SSA (Garcia-Marques et al., 2019) (Figure 3G). *ap*-driven expression of Cas9 in combination with *U6*-gRNA#2 resulted in robust dorsal activation of the Actin5C-CAAX-mCher(#2)rry CaSSA reporter (Fig 3H). Consistent with the observations made when switching of SEED cassettes (Figure 3C, D), numerous clones expressing mCherry:CAAX localize outside the dorsal compartment (Figure 3H). The use of *UAS*-driven gRNAs resulted in a drastic reduction of off-target labeling outside the dorsal compartment (Figure 3H’). Both *ihog*^sfGFP-SEED^ and Actin5C-CAAX-mCher(#2)rry were then combined to trigger simultaneous cell labelling (via CaSSa) and gene labelling (via SEED) by expressing Cas9 and gRNAs under *ap*-Gal4 control. The co-labeling of Ihog:sfGFP and cell membranes (mCherry:CAAX) covered the dorsal compartment (Figure 3I), demonstrating the compatibility of the two methods.

One of the limitations of combining SEED cassettes with CaSSA reporters is the presence of the 3xP3-FP marker. The 3xP3 enhancer drives expression in numerous tissues throughout development, especially in the brain and the eyes (Figure 1D) (Horn, Jaunich, & Wimmer, 2000), impeding simultaneous cell-labeling and endogenous gene tagging in these tissues. To overcome this limitation, we designed a SEED cassette (*SEED-LoxP*) that permits Cre-mediated removal of the 3xP3 marker prior to harvesting (Supplementary figure 4D, E). Using this approach, we generated an *ihog*^sfGFP-SEED-LoxP^ allele. The knock-in efficiency using this donor was similar to that obtained for *ihog*^sfGFP-SEED^ (Supplementary figure 4F).

In order to test the capacity of Cas9 to direct endogenous labeling in other tissues, we activated the *ihog*^sfGFP-SEED^ cassette using the neuronal driver lines *vGlut*-Cas9 and *elav*-Cas9 (Supplementary figure 4A, B, B’). In both cases, labeling was easily detected in neural somas along the ventral nerval chord (VNC) of third instar larvae, indicating the feasibility of using this approach in the brain. Interestingly, activation of *ihog*^sfGFP-SEED^ using elav*-*Cas9 resulted in large Ihog:sfGFP clones in the wing disc (Supplementary figure 4C), due to the transient activity of the *elav* promoter at low levels in imaginal discs (Casas-Tintó, Arnés, & Ferrús, 2017).

Together, these results demonstrate efficient tissue-specific gene labeling triggered by CRISPR/Cas technology. A new CRISPR application that can be easily combined with other Cas9-triggered applications.

### Conditional rescue of mutant SEED alleles

When SEED cassettes are inserted in the 5’ region of a gene, translation will be prematurely interrupted by a stop codon after the first repeat of the cassette, resulting in the generation of a mutant allele. We envisioned that these mutants could be rescued by conditional expression of Cas9 and harvesting gRNAs. To test this possibility, we targeted the *shf*^sfGFP-SEED^ allele produced upon sfGFP-SEED cassette knock-in in the *shf* locus (Figure 2H, Figure 4A). *shf* is expressed in the lateral regions of the wing and encodes a highly diffusible protein strictly required for Hh dispersal (Glise et al., 2005; Gorfinkiel, Sierra, Callejo, Ibanez, & Guerrero, 2005). Cassette insertion is predicted to generate a truncated protein only preserving the first 60 amino acids of Shf (Figure 4A). In accordance, *shf*^sfGFP-SEED^ hemizygous males phenocopied the previously described *shf* null alleles, displaying a reduction of the 3-4 intervein wing territory (Figure 4B’) (Glise et al., 2005; Gorfinkiel et al., 2005; Lindsley & Zimm, 1992), reduced and rough eyes (Figure 4I, Supplementary Figure 5F) (Bateman, 1950; Lindsley & Zimm, 1992), and loss of scutellar bristles (quantified in Figure 4H) (Lindsley & Zimm, 1992). As expected, no GFP fluorescence was detected in *shf*^sfGFP-SEED^ wing discs (Figure 4B). Hh signaling range, revealed by immunostaining of target genes Ptc and full-length Cubitus interrupturs (Ci^155^), was constrained to the first anterior cell row in contact with the posterior compartment (Supplementary Figure 5A, A’).

**Figure 4.**
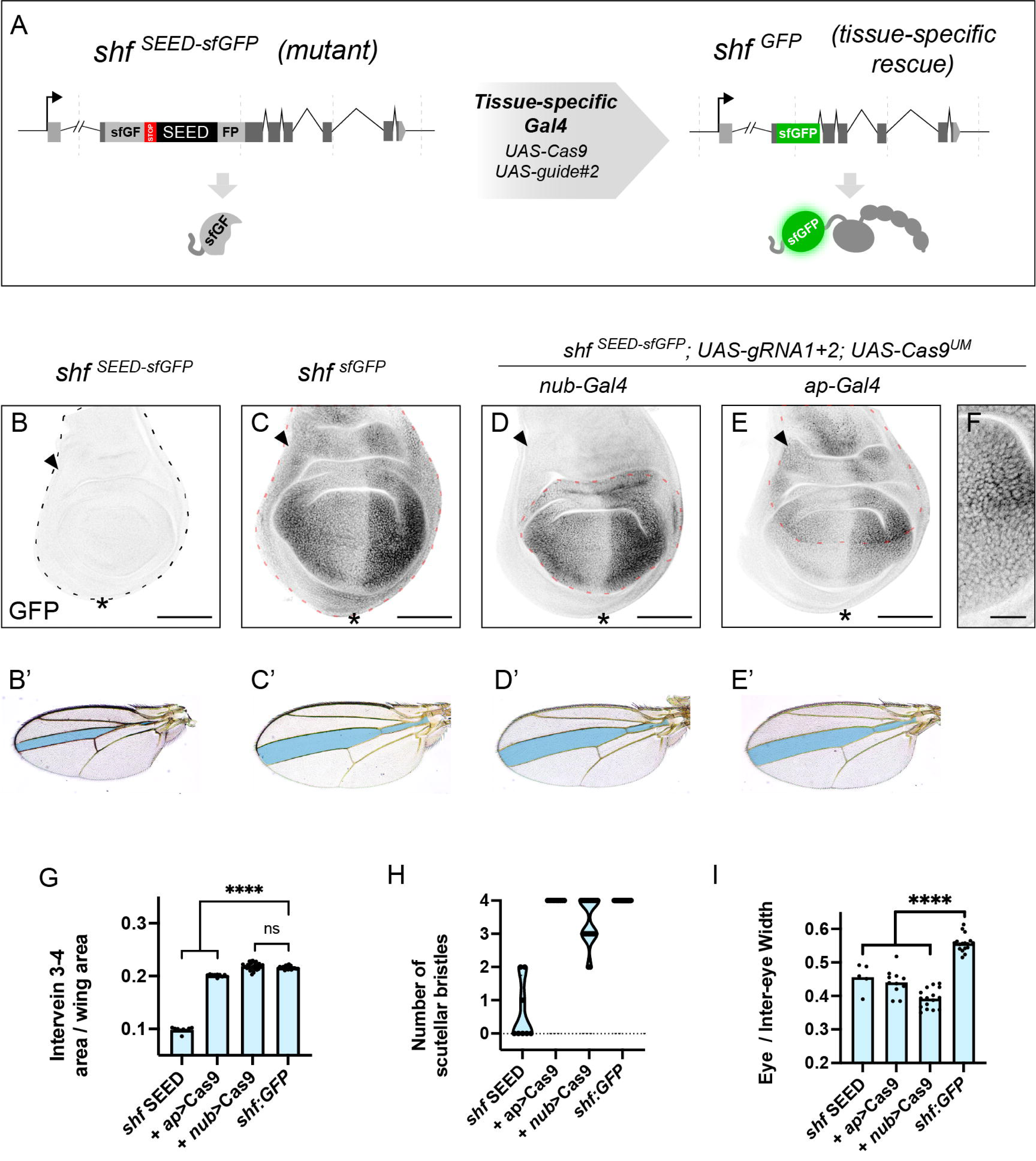
Tissue-specific rescue of mutant alleles. **A.** Scheme of the rescue strategy. N-terminal insertion of the sfGFP-SEED cassette results in a premature stop cassette after the first repeat, leading to a truncated protein containing only the first 60 amino acids of Shf and a non-fluorescent truncated sfGFP. Tissue-specific expression of Cas9 and harvesting gRNAs harvests the SEED cassette resulting in endogenous tissue-specific expression of Shf tagged with GFP. **B.** GFP fluorescence of a *shf^S^*^EED-sfGFP^ control discs. **C.** GFP fluorescence of a *shf^s^*^fGFP^ wing disc (after germline excision of the SEED cassette). **D.** Tissue-specific rescue of the *shf^S^*^EED-sfGFP^ allele within the *nub* expression domain (delimited by the red dotted line). Notice the rescue of GFP levels within the pouch. Arrowhead indicates the reduced levels in the proximal hinge region (compare with panel C). Asterisk indicates the reduced levels in the ventral hinge (compare with panel C). **E.** Tissue-specific rescue of the *shf^S^*^EED-sfGFP^ allele within the *ap* expression domain (delimited by the red dotted line). GFP signal is higher in the dorsal compartment, with reduced levels in the ventral pouch. Arrowhead indicates the presence of GFP in the proximal hinge region (compare with panel C and D). Asterisk indicates the reduced levels in the ventral hinge (compare with panel C and D). **F.** Detail of sfGFP:Shf distribution in the posterior wing pouch from the wing disc of panel E. **B’-E’.** Adult wings of the genotypes depicted in panels B to E. The adult wing of *shf^S^*^EED-sfGFP^ presents a phenotype that mimics that of null mutants (Glise et al., 2005; Gorfinkiel et al., 2005), with reduced 3-4 intervein territory (labelled in blue). **G.** Quantification of intervein 3-4 relative size. *shf^S^*^EED-sfGFP^ n=9, *shf^s^*^fGFP^(*ap*) n=10, *shf^s^*^fGFP^(*nub*) n=28, *shf^s^*^fGFP^ n=17. p value<0.0001. **H.** Number of scutellar bristles in the different backgrounds. *shf^S^*^EED-sfGFP^ n=8, *shf^s^*^fGFP^(*ap*) n=10, *shf^s^*^fGFP^(*nub*) n=20, *shf^s^*^fGFP^ n=26. **I.** Quantification of the relative eye size in the different backgrounds, calculated as the total eye with divided by the inter-eye region. *shf^S^*^EED-sfGFP^ n=5, *shf^s^*^fGFP^(*ap*) n=11, *shf^s^*^fGFP^(*nub*) n=16, *shf^s^*^fGFP^ n=17. p value>0.0001. Scale bars: B, C, D, E: 100μm; F: 25 μm

Germline harvesting (excision) of the SEED cassette resulted in GFP detection in the wing disc (Figure 4C), in a pattern that mimicked previously published Shf antibody stainings (Glise et al., 2005). Hh signaling displayed normal range, Ptc being detected in the first four rows of anterior cells (Supplementary Figure 5B). The long-range target Ci^155^ was detected in a graded manner, with its characteristic repression in the first cells of the anterior compartment (Supplementary Figure 5B’). The phenotype of the adult wing was also rescued (Figure 4C’), as well as the scutellum and the eye defects (Figure 4H, I, Supplementary Figure 5G, G’). Together these results confirmed that the sfGFP:Shf protein is functional and, under these experimental conditions, equivalent to untagged Shf.

Spatio-temporal control of *shf*^sfGFP-SEED^ rescue was then attempted by providing Cas9 and harvesting gRNAs in somatic cells, under the control of *nub*-Gal4. *nub*-driven rescue resulted in a normal sfGFP:Shf signal in the wing pouch (Figure 4D). Despite of the high diffusivity of Shf, GFP levels were lower in the hinge and notum than those of *shf*^sfGFP^ controls, suggesting that those structures did not produce the GFP-tagged protein. Hh signaling range was fully rescued in the wing pouch (Supplementary Figure 5C, C’). Accordingly, the relative size of the intervein 3-4 was identical to *shf*^sfGFP^ wings (Figure 4D’, G). The number of scutellar bristles, dependent on Hh signaling in the notum, was only partially rescued (Figure 4H). Consistent with the restricted rescue of *shf*, eyes still displayed mutant phenotype (Figure 4I, Supplementary Figure 5H, H’), most likely due to the lack of expression of *nub*-Gal4 in the imaginal eye disc.

The *shf*^sfGFP-SEED^ allele was then rescued by the expression of Cas9 and harvesting gRNAs under the dorsal driver *ap*-Gal4. Consistent with localized protein production, sfGFP:Shf was predominantly detected in the dorsal compartment (Figure 4E, F), present at lower levels on ventral cells due to extracellular dispersal. Hh signaling was partially rescued in the wing pouch, with reduced range in the ventral compartment, far from the dorso-ventral boundary (Supplementary Figure 5D, D’, E). Accordingly, the relative size of the intervein 3-4 was not totally rescued (Figure 4E’, G). In contrast to the *nub*-driven rescue, the dorsal hinge and the notum of *ap*-rescued animals presented normal levels of sfGFP:Shf (Figure 4E). In addition, the number of scutellar bristles was fully rescued (Figure 4H). Eyes still displayed the mutant phenotype (Figure 4I, Supplementary Figure 5I, I’).

Collectively, these results demonstrate that SEED cassettes can be used to achieve spatio-temporal control of protein expression, via CRISPR-triggered mutant rescue.

### Visualization of ALFA-tagged knock-ins by a conditionally-stable ALFA Nanobody

SEED cassettes permit tissue-specific labeling with fluorescent proteins (Figure 3). However, SEED cassettes bearing short peptide tags produce a tagged protein even before harvesting, and thus, cannot be conditionally activated. As an alternative to visualize these knock-ins, we propose indirect labeling via chromobodies. Chromobodies are genetically-encoded fusions between a protein binder and a fluorescent protein. Several chromobodies have been developed for Drosophila (Kamiyama et al., 2015; Vigano et al., 2021; Xu et al., 2022). However, their implementation to detect endogenously tagged proteins is challenging, due to the high background produced by the unbound chromobodies, that accumulates in the cytoplasm when its expression is higher than the target protein (reviewed in Aguilar, Vigano, et al., 2019).

In order to develop useful chromobodies, we chose ALFANb{Götzke, 2019 #502}, given its high affinity for ALFA peptide and its stability *in vivo.* ALFANb was fused to mCherry and expressed in wing imaginal discs using *en-*Gal4, both in the absence (Figure 5B upper row) and the presence of Sqh:ALFA:HA (Figure 5B lower row). *sqh* encodes the Drosophila homolog of non-muscular myosin II, a protein that localizes in the cellular cortex. In the presence and in the absence of Sqh:ALFA:HA, the ALFA-chromobody totally filled the cytoplasm, rendering the visualization of Sqh impossible due to the high background (Figure 5B, first column panels). While reduction of expression levels is possible via titration of the Gal4/UAS expression system, this would be incompatible with many experimental setups.

**Figure 5.**
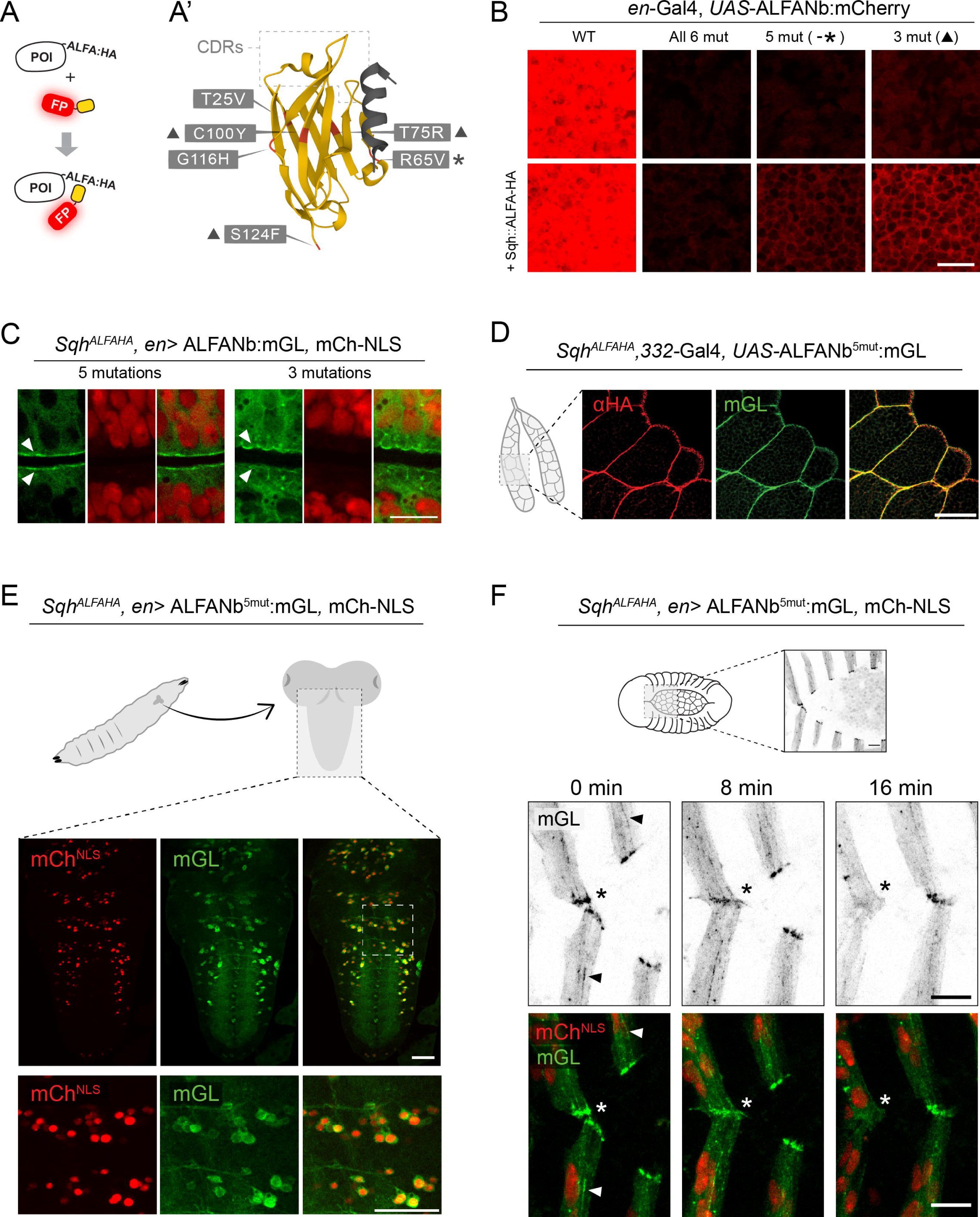
Protein visualization via conditionally-stable ALFA nanobodies. **A.** Scheme of the chromobody concept. Upon co-expression of an ALFA-tagged protein of interest (POI) and the anti-ALFA chromobody (fusion of the ALFANb and a fluorescent protein (FP)), the ALFA nanobody would be recognize the ALFA epitope, permitting the fluorescent visualization of the POI via the FP. **A’.** Structure of the ALFANb (yellow) bound to the ALFA peptide (grey) (PBD:2I2G, (Götzke et al., 2019). Substitutions introduced to make it unstable are highlighted in red. Note that R65V is located on the surface contacting ALFA. The 6-mutation version includes all 6 substitutions. The 5-mutation includes all but R65V (marked with an asterisk). The 3-mutation version includes only T75R, C100Y and S124F (marked with a triangle). **B.** Fluorescence of the different versions of the ALFANb fused to mCherry using *en-*Gal4 in absence (upper row panels) or presence (lower row panels) of endogenously tagged Sqh:ALFA:HA. ALFANb-mCh (first column) presents similar levels in both presence and absence of Sqh:ALFA:HA, filling the cytoplasm. ALFANb^6mut^-mCh (second column) is highly destabilized in absence of Sqh:ALFA:HA, but fails to localize cortically in its presence. ALFANb^5mut^-mCh (third column) is still destabilized in Sqh:ALFA:HA absence, localizing in the cell cortex in its presence. ALFANb^3mut^-mCh (fourth column) results in higher levels than ALFANb^5mut^-mCh (third column) in both absence and presence of tagged Sqh. **C.** Detail view of the wing disc fold upon expression of ALFANb^5mut^-mGL and ALFANb^3mut^-mGL in the presence of Sqh:ALFA:HA In green, ALFANb-mGL fusions. In red, mCherry:NLS. ALFANb^5mut^-mGL resulted in enrichment in the apical area, limiting with the fold’s lumen (arrowheads). Only low GFP levels can be detected in the nucleus. ALFANb^3mut^-mGL results in apical localization. Cytosolic and nuclear levels are higher than those of ALFANb^5mut^-mGL. **D.** Confocal images of ALFANb^5mut^-mGL expressed in the salivary glands in the presence of Sqh:ALFA:HA. Both anti-HA signal (red) and mGL fluorescence (green) can be detected colocalizing in the periphery of the cells. Some cytoplasmic signal is detected for both HA and mGL. In the right panel, merge of both mGL and anti-HA signals. **E.** Confocal images of ALFANb^5mut^-mGL expression in the brain of Sqh^ALFA:HA^ larvae using *en*-Gal4. Nuclei are labelled via UAS-mCherry:NLS (red panels). mGL signal (green panels) can be detected in the cell body, mostly excluded from the nucleus, and in cellular extensions. Right panel contains merged signal of both red and green panels. Lower row of panels portrays a magnification of the inset labeled in the upper-right panel. **F.** Stills from three time-points of Supplementary video 2, portraying epidermal segment fusion during dorsal closure in Sqh^ALFA:HA^ embryos. *en*-Gal4 is used to drive ALFANb^5mut^-mGL and mCherry:NLS in the posterior segmental compartments of the epidermis. Upper row depicts mGL signal alone (in black). Lower panels depict a merged composition of both mGL and mCherry signals. mGL signal is detected in bundles and foci at the cortex of the cells in the posterior segmental compartments (arrowheads) and is especially enriched in the leading-edge cells forming the actomyosin cable. Upon leading edge cell contact formation (0min) and fusion of the epidermal segments (8min) the Sqh foci located in the leading edge, dissolve in the cytoplasm (16min) Asterisk depicts the different stages of fusion, from recognition (0 min, left panel) to fusion (8 min, middle panel) and removal of the Sqh foci (16min, right panel). Scalebars: C: 10μm, D: 100μm, E: 50μm, F: 10μm

Thus, we decided to engineer the nanobody to be degraded in the absence of its target. To that end, we introduced the six mutations proposed to result in conditional stability of the anti-GFP nanobody (J. C. Tang et al., 2016) (Figure 5A). These mutations destabilized the chromobody (Figure 5B second column panels, compared to first column ones), but did not result in cortical accumulation when co-expressed with Sqh:ALFA:HA (Figure 5B second column panels, upper row vs lower row), most likely due to loss of target affinity. The ALFANb interacts with the ALFA peptide partially via its complementarity-determining regions (CDRs) but also with the lateral surface of the nanobody core ((Götzke et al., 2019), Figure 5A). One of the mutations introduced (R65V) was located on the interacting surface with the ALFA peptide. The ALFA-chromobody bearing the other five mutations was still largely unstable when expressed alone, but localized cortically if co-expressed with Sqh:ALFA-HA, indicative of Sqh binding (Figure 5B third column panels). We also generated an ALFA-chromobody mutated in the three core positions (T75R, C100Y and S124F) shown to have the strongest impact on conditional stability, and proposed by Cepko and colleagues to be the most transferrable across nanobodies (J. C. Tang et al., 2016). This form was slightly less destabilized than the five-mutation version (Fig 5B, compare third and fourth column panels), as expected from previous results (J. C. Tang et al., 2016). Likewise, it resulted in higher signal when co-expressed with Sqh:ALFA:HA (Figure 5B fourth column, second row panel).

In order to obtain a chromobody compatible with high resolution imaging, we generated fusions of both ALFANb^5mut^ and ALFANb^3mut^ with the much brighter fluorescent protein mGreenLantern (mGL) {Campbell Benjamin, 2020 #774}. Expression of either form in the wing disc using the *en*-Gal4 driver resulted in cortical localization, highly enriched in the apical region (Figure 5C). Nevertheless, ALFANb^3mut^-mGL presented poorer signal-to-noise ratio when compared to ALFANb^5mut^-mGL, as revealed by the higher fluorescence levels in the nucleus, were Sqh is normally absent (Figure 5C, arrowheads). Thus, we decided to employ ALFANb^5mut^-mGL for all further experiments. To demonstrate the extent to which the ALFANb^5mut^-mGL reflects Sqh localization, *332.3*-Gal4{Wodarz, 1995 #1164} was used to drive its expression in the salivary glands. In the large cells of this tissue, the cortical localization of ALFANb^5mut^-mGL was identical to that of Sqh:ALFA:HA, as revealed by anti-HA immunostaining (Figure 5D). Both anti-HA and ALFANb^5mut^-mGL fluorescence were present at low levels in the cytoplasm.

One of the tissues in which conditional protein labeling would be most useful is the brain. This tissue is highly packed and its cells have intricate tridimensional morphologies, complicating the study of protein localization. In the larval brain, expression of ALFANb^5mut^-mGL using *en*-Gal4 in a *sqh*^ALFA:HA^ background resulted in the labelling of myosin-rich structures in the *en*-positive cells. Sqh:ALFA:HA was detected in the cell body, surrounding the nucleus, and labeled neurites that cross the ventral nerve cord (VNC) (Figure 5E).

### Live-imaging of ALFA-tagged knock-ins

Chromobodies have become versatile tools to label endogenous proteins and study their dynamics *in vivo* (Panza et al., 2015). Actomyosin dynamics, of which the motor protein Sqh is a core component, govern several morphogenetic processes (Munjal & Lecuit, 2014). Among them, embryonic dorsal closure is often used as a model of epidermal morphogenesis and wound healing (Kiehart, Crawford, Aristotelous, Venakides, & Edwards, 2017). During dorsal closure, the two lateral epidermal cell sheets of the embryo move up and converge at the dorsal-midline to seal the epithelial gap, initially occupied by the amnioserosa. The leading edge cells of each epidermal sheet form a supracellular actomyosin cable that surrounds the dorsal opening, contributing to closure dynamics and preventing morphogenetic scarring (Ducuing & Vincent, 2016). Dorsal closure completes by the zipping of the two epidermal cell sheets, resulting in the perfect matching of the epidermal segments. This process requires filopodial and lamellipodial protrusions emerging from the leading edge cells of the epidermis (Jacinto et al., 2000). Expression of ALFANb^5mut^-mGL via *en*-Gal4 in a *sqh*^ALFA:HA^ background resulted in Sqh labeling in the posterior segmental compartments of the epidermis (Figure 5F, asterisk, Supplementary Movie 1). The mGL signal was outlining the epidermal cells (Figure 5F, arrowhead, Supplementary Movie 1) and was strongly enriched in distinct foci at the actomyosin cable, consistent with previously reported Sqh:GFP dynamics (Franke, Montague, & Kiehart, 2005). We further observed the mGL signal as part of cellular protrusions in leading edge cells (Figure 5F, Supplementary Movie 1). Upon fusion of the epidermal segments at the dorsal midline, the mGL signal disappeared (Figure 5F, 16min time-point, Supplementary movie 2). In summary, our observations demonstrate that ALFANb^5mut^-mGL can faithfully capture the localization and dynamics of Sqh *in vivo*.

### Simultaneous manipulation and visualization of short-peptide knock-ins

While protein visualization is critical for many studies, direct protein manipulation is fundamental to assess functional roles. We have previously proposed deGradFP, a method to target GFP-tagged proteins for degradation. deGradFP is a fusion of an anti-GFP nanobody to a F-box, which recruits the ubiquitination machinery, subsequently leading to proteasomal degradation of the GFP-tagged protein (E. Caussinus, Kanca, & Affolter, 2011). We tested whether the anti-GFP nanobody could be substituted by ALFANb to target ALFA-tagged proteins. We named this tool degradALFA. Following deGradFP’s design, the F-box of the protein Slmb was chosen for the functionalization (Figure 6A). Expression of degradALFA in the posterior compartment using *en*-Gal4 in a *sqh*^ALFA:HA^ homozygous background resulted in embryonic lethality (data not shown), consistent with previous reports using deGradFP (Ochoa-Espinosa, Harmansa, Caussinus, & Affolter, 2017). In order to investigate the degree of Sqh degradation in larval stages, we rescued embryonic lethality by maternally providing untagged Sqh. These animals reached larval stages. Consistent with protein degradation, the anti-HA signal was dramatically reduced in posterior wing disc cells (Figure 6B). According to previous results (Emmanuel Caussinus & Affolter, 2016; Ochoa-Espinosa et al., 2017; Pasakarnis, Frei, Caussinus, Affolter, & Brunner, 2016), punctated non-degraded accumulations were detected in the basal part of the disc epithelium. Immunostaining using anti-ALFA failed to detect Sqh in posterior cells, further indicating Sqh degradation or that the ALFA epitope is being covered by the nanobody of degradALFA (Figure 6C). The size and shape of the affected posterior compartment was also perturbed in respect to the anterior. Recently, we have proposed the degradation of HA-tagged proteins using degradHA, a functionalized anti-HA single-chain antibody (scFv) (Vigano et al., 2021) (Figure 6D). We evaluated whether Sqh could be targeted by this method and to which extent Sqh was degraded. In contrast to degradALFA, *en*-Gal4-driven degradHA in a *sqh*^ALFA:HA^ background did not result in embryonic lethality. However, consistent with degradation, the wing discs of these animals displayed reduced anti-ALFA immunostaining in the posterior compartment (Figure 6E). In addition, wing disc cells also presented the characteristic Sqh aggregates that are seen upon the expression of degradALFA and deGradFP (Figure 6E, (E. Caussinus et al., 2011)). Anti-HA immunostaining result in little signal in the posterior compartment (Figure 6F). These results indicate that degradHA is also able to trigger Sqh:ALFA:HA degradation, but most likely to a lesser extent than degradALFA.

**Figure 6.**
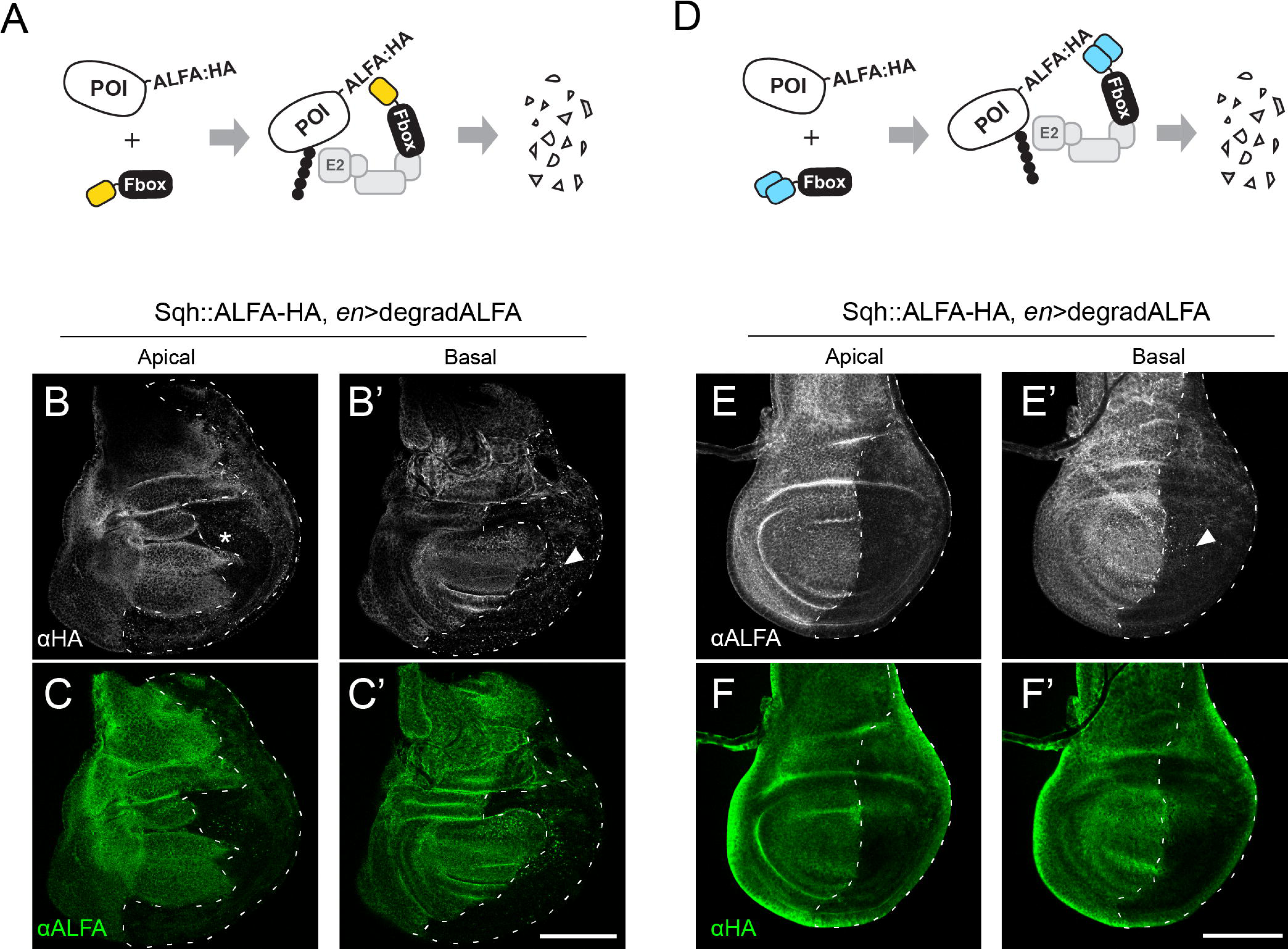
Simultaneous degradation and visualization of double-tagged knock-ins. **A.** Scheme of ALFANb-mediated degradation. The ALFANb (in yellow) is fused to the F-box domain contained in the N-terminal part of Slmb (in black). Upon binding of an ALFA-tagged peptide, the F-box recruits the polyubiquitination machinery, targeting the POI to degradation. **B-C.** Single Z stacks of wing discs expressing degradALFA in the posterior compartment (delimited by a dotted line) using the *en-*Gal4 driver. In grey, anti-HA immunostaining (B and B’). In green, antiALFA immunostaining (C and C’). Notice the not-straight shape of the AP boundary in the pouch. An apical (B, C) and a basal (B’, C’) planes are shown. In white, an anti-HA staining reveals strong reduction in Sqh levels. Dotted signal can be observed, especially in the basal plane (arrowheads). The posterior compartment presents interruption of the hinge folds (asterisk). In green, immunostaining using anti-ALFANb:488. Posterior compartment presents also a strong reduction of ALFANb:488 levels compared to the anterior compartment. This signal reduction is probably exacerbated by the masking of ALFA peptide by the ALFANb of degradALFA. **D.** Scheme of degradHA-mediated protein targeting. The HA scFv (in blue) is fused to the F-box domain (in black) contained in the N-terminal part of Slmb. Upon binding of an HA-tagged peptide, the F-box recruits the polyubiquitination machinery, targeting the POI to degradation. **E-F.** Wing discs expressing degradALFA in the posterior compartment (delimited by a dotted line) using the *en-*Gal4 driver. In grey, anti-ALFA immunostaining (E and E’). In green, antiHA immunostaining (F and F’). An apical (E and F) and a basal (E’ and F’) planes are shown. In white, an anti-ALFA immunostaining reveals strong reduction in Sqh levels. Dotted signal can be observed in the basal plane (arrowheads). In green, anti-HA immunostaining. Posterior compartment presents strongly decreased anti-HA levels compared to anterior compartment, further indicating that Sqh levels are reduced. Scalebars: 100μm.

Protein degradation is a valuable tool for assessing the role of proteins. Nevertheless, in many cases, degradation is not the most suitable strategy, due to the nature of the targeted protein or the intended manipulation. Many different protein binder-based approaches are currently in use in developmental biology (Aguilar, Matsuda, Vigano, & Affolter, 2019; Bieli et al., 2016). In these lines, we foresaw a new method to prevent secretion of proteins tagged with ALFA in the cytoplasmic domain. For this purpose, we fused the ALFA nanobody to the cytoplasmic tail of CD8, used as a transmembrane scaffold in other nanobody-based toolkits (Stefan Harmansa, Alborelli, Bieli, Caussinus, & Affolter, 2017; S. Harmansa, Hamaratoglu, Affolter, & Caussinus, 2015). To reinforce ER retention, we added the retention signal KKXX, present in KDEL receptors, to its C -terminus (Supplementary Figure 6A). For visualization purposes, CD8 was tagged in the N-terminus with mCherry. We refer to this construct as ALFA-Trap^ER^. We hypothesized that co-expression of ALFA-Trap^ER^ together with a secreted protein tagged in the cytoplasmic tail, would result in nanobody binding and retention via the KKXX signal. Expression of ALFA-Trap^ER^ using the wing pouch-driver *pdm2-*Gal4 in a *ptc*^ALFA:HA^ background resulted in strong build-up of anti-HA signal compared to controls (Supplementary Figure 6B). This data suggest that Ptc is retained in those cells, likely resulting in Hh signaling up-regulation and thus more Ptc being produced. Given that all Ptc is tagged with ALFA:HA, this would result in the further accumulation of Ptc:ALFA:HA in the ER and thus the observed build-up of anti-HA signal.

In sum, our data demonstrate the versatility of gene knock-ins with short peptide tags in tandem. Together with the novel nanobody-based toolbox, based on the ALFANb, this knock-ins permit tissue-specific control over the visualization and/or manipulation of their protein products.

## Discussion

Here, we have developed a novel strategy, called SEED/Harvest, to generate knock-ins in Drosophila. Similar to other knock-in technologies, SEED relies on the HDR pathway to insert exogenous DNA in the targeted locus. In contrast to other alternatives, CRISPR is also responsible in an SSA-dependent process for the removal of the marker used for screening. Its sole dependency on CRISPR/Cas makes SEED a powerful alternative to current methods and opens a series of experimental possibilities. We have demonstrated the versatility of the method by tagging relevant genes of the Hh pathway, thus generating a valuable toolbox. We now use SEED to routinely generate knock-ins and validated its use in several loci, obtaining robust efficiency of insertion and harvesting (Bauer, Aguilar, Wharton, Matsuda, & Affolter, 2023; Mesrouze et al., 2022).

In contrast to most methods, SEED permits scarless tagging of the targeted locus. This property is fundamental for many purposes, such as internal tagging of proteins or the generation of point mutants. In these cases, scars could disrupt the protein sequence in unpredictable ways. Scarless tagging can also be achieved when using templates that do not contain a selectable marker for screening (Port et al., 2014a). These templates permit the generation of scarless knock-ins in one step. However, they require the screening for successful editing to be performed via PCR, as most of the tagged proteins are hardly visible with fluorescence binoculars. While this approach can be used to generate a few knock-ins at a time, scaling up their generation seems unfeasible. Moreover, efficiency of insertion varies among loci. In this study, the insertion in the *shf* locus led to only two founder parents out of 30 fertile F0 individuals (Figure 2H). In that case, PCR screening would have been likely to fail.

To date, PBac technologies also permit rapid screening and scarless marker removal in Drosophila (https://flycrispr.org/ (Nyberg et al., 2020). This technology has demonstrated to be efficient in many loci. SEED and PBac technologies differ in several ways. The first difference is the dependency on a TTAA site. These sites are abundant in introns and other non-coding regions but less frequent in exonic sequences. While TTAA sites can be added via silent mutations or included in linker sequences, they generally complicate the cloning or result in insertion of amino acids rarely used in linkers (Chen, Zaro, & Shen, 2013). The second difference strides on the machinery needed for marker removal. The dependency on PBac transposase is not a problem in Drosophila, where many genetic tools have already been established. Nevertheless, when applying these technologies in other organisms, the SEED/Harvest’s dependence only on CRISPR/Cas turns advantageous. Moreover, expression of transposase entails the risk of transposon insertion in other loci, with potentially unpredictable consequences. Finally, SEED/Harvest technology can be used in combination with other CRISPR-based, tissue-specific tools, opening new experimental avenues.

The SEED approach also presents inherent limitations in certain set-ups. The most obvious is the integration of highly repeated sequences. In these cases, SSA-mediated removal of the selectable marker is likely to produce many different outcomes; in such cases, SEED/Harvest is not the most suitable alternative.

Another popular approach for gene editing in Drosophila involves the CRISPR-mediated insertion of *attP* sites (Baena-Lopez et al., 2013; Huang, Zhou, Dong, Watson, & Hong, 2009). *attP* provides a landing site into which *attB*-containing DNA can be inserted. These approaches are very modular, and permit, once the *attP* insertion has been generated, robust integration of different genetic variants. On the other hand, these approaches involve several rounds of insertion and marker removal, with the concomitant increase in time. In Drosophila, these methodologies often involve several rounds of injection and can take more than 10 generations, complicating its implementation in vertebrates. SEED/Harvest overcomes these limitations, permitting fast and robust generation of knock-ins. These characteristics make it a potential candidate methodology for scarless gene editing in vertebrates.

For many laboratories, one of the most time-consuming steps to generate knock-ins is the cloning of donor vectors. Amplification and cloning of long homology arms is often challenging. One of the main advantages of SEED/Harvest is the simplicity by which donor templates are generated. In this study, we provide reagents to tag genes with common tags used for protein purification and visualization, as well as fluorescent proteins (Figure 1E, Supplementary Figure 1F). In addition to these tools, we have adapted the SEED/Harvest technology to be used with short homology arms (<200bp). Small homology arms had been proven successful for integration of other donors (Kanca et al., 2019), and we did not observe a major decrease in integration efficacy respect long homology arms (Figure 2). These rates are higher than those previously described using short homology arms (Kanca et al., 2019). While it is unclear whether these differences are due to the SEED/Harvest strategy or to the injection procedure, they highlight the potential of the strategy.

Donors with short homology arms require vector synthesis by a commercial provider and a one-day cloning step. For most approaches, this strategy will be satisfactory with regard to time and costs. Nevertheless, we envisioned a way to optimize both. Inspired by previous reports (Wierson et al., 2020), we employed even shorter homology arms to generate donors. This approach permitted one day cloning, without the delay that synthesis implies, while maintaining a considerable efficiency compared to 185bp homology arms (Figure 2L, M). This is the first time this strategy is used in Drosophila and it questions the requirement of the 500bp-1500bp length of homology arms in most CRISPR approaches.

A distinctive feature of SEED cassettes is the possibility to remove the marker in somatic cells, and therefore induce endogenous tagging. In fact, these insertion cassettes are by-products of the generation of the scarless knock-ins. Conditional gene labeling has been achieved in the past via careful locus engineering (Alexandre et al., 2014). While yielding very robust results, the applicability of this approach might be limited in other organisms, given the many generations required to create these alleles. Recently, several tools based on artificial intron tagging have been proposed to mediate conditional gene labeling via restricted Flippase expression (Fendl, Vieira, & Borst, 2020; Nagarkar-Jaiswal, Manivannan, Zuo, & Bellen, 2017; Williams, Shearin, & Stowers, 2019). These studies provide a valuable asset for many genes where intronic tagging is possible. However, many proteins require internal, N-or C-terminal tagging, which would not be suitable for such a design. In turn, SEED permits gene labeling upon CRISPR-mediated DSB. In the past years, tissue-specific expression of Cas9 has been proposed to trigger numerous events, from gene knock-out (Koreman et al., 2021; Meltzer et al., 2019; Port, Chen, Lee, & Bullock, 2014b) to cell-(Koreman et al., 2021) and membrane-labeling (Garcia-Marques et al., 2019). SEED/Harvest can be easily combined with these approaches to design increasingly complex experiments while reducing the number of tools needed.

In many cases, the generation of knock-ins aims for the tagging of a protein with a short tag or a fluorescent protein. In this study, we propose the tagging with small protein tags in tandem. Recently, we and others have proposed the *in vivo* manipulation of proteins via these short tags, demonstrating that one copy is enough for efficient manipulation (Vigano et al., 2021; Xu et al., 2022), and a second, different small tag is essential for antibody staining upon manipulation. To complement the genetic tools presented in this study, we developed a toolbox to visualize and manipulate proteins via the ALFA nanobody (Götzke et al., 2019).

Visualization of proteins through chromobodies depends on a tight balance between the levels of the target protein and the chromobody itself. It has been proposed that many nanobodies are inherently destabilized in absence of antigen (Keller et al., 2018). We found this not to be the case for the ALFA nanobody (Figure 4B). To generate a useful chromobody, we engineered the ALFA nanobody such that it would be degraded unless bound to the ALFA tag. We grafted both the mutations responsible of the conditionally-stable behavior of the anti-GFP nanobody dGBP as well as the three major mutations proposed to be the most transferrable across nanobodies (J. C. T. Tang et al., 2016). Testing certain combinations of these mutations permitted us the isolation of a functional destabilized chromobody (Figure 5). The destabilized ALFA nanobody that we present here is the first conditionally unstable nanobody detecting a short tag, with many potential applications.

Together, these tools permit the robust, affordable and rapid generation of knock-ins, as well as conditional gene tagging, protein visualization and protein manipulation.

## Material and methods

### Immunohistochemistry and imaging

3^rd^ instar larvae of the indicated genotype were dissected in cold PBS (PH 7.2, Gibco™ (PN: 20012019)), immediately followed by fixation at room temperature for 30 min using a 4% paraformaldehyde solution (Electron Microscopy Sciences, (PN:15714)) in PBS. After thorough washing with PBS, samples were directly mounted or subjected to immunohistochemistry. For immunohistochemistry, samples were permeabilized in PBST (PBS + 0,3% Triton X-100 (Sigma PN:1002194227)) for 30 min, followed by 1h incubation in blocking solution (5% Normal Goat Serum (Abcam, ab7481) in PBST). Primary antibodies were diluted in blocking solution and used to incubate samples overnight at 4 degrees. The next day, samples were washed with PBST for 3×15min and incubated for 2 hours in secondary antibody diluted in blocking solution. A final 3×15min washing in PBST and 2×15min washing in PBS was performed before mounting. Samples were gently rotated during fixation and immunostaining. For mounting, samples were placed in Vectashield® with (PN: H-1200) or without DAPI (PN: H-1000). Tissues where then separated from the larval cuticle on a glass slide. A cover-slide was placed on top and sealed with nail polish. Preparations were imaged using LSM880 confocal and analyzed by OMERO and imageJ.

### Antibodies used in this study

The following antibodies were used in this study: anti-HA 1/200 (3F10 clone, sigma), anti-Ptc 1/150 (Apa1.3, isolated from hybridoma cells (Capdevila et al., 1994)), anti-Ci^155^ (2A1, Developmental Studies Hybridoma Bank), 488 conjugated anti-ALFA 1/500 (clone 1G5, Nanotag Biotech.), Alexa fluor 568 anti-Rat (Invitrogen, A11077),.

### Single fly PCR

Flies were anesthetized with CO2 and placed in PCR tubes on ice. Samples were homogenized using a plastic tip in 30μL of squishing buffer (10mM Tris-HCl, 2.5mM EDTA, 25mM NaCl, 200ng/mL Proteinase K). Following, samples were incubated at 37 degrees for 30min. Proteinase K was subsequently inactivated by heating at 95 degrees for 5min. 1μL was used for downstream PCRs.

### Generation of donor vectors

Most 185bp-long plasmids generated in this study where cloned prior the adoption of Golden Gate, which we incorporated later to reduce the cloning time e improve the modularity (for description of this methodology see next section of material and methods). The cassettes of *ihog*, *boi*, *shf*, and *en*, were cloned as follows: A plasmid containing the guideRNAs targets, homology arms and two tandem BsaI target sites was synthesized by Genewiz. To facilitate homology directed assembly (Gibson or NEB Hi Fi assembly), 30bp homology with the preferred SEED cassettes were included at each site of the BsaI sites, in frame with the selected SEED cassette. All the synthesized fragments are included in the Extended materials. When necessary, silent mutation were introduced to avoid cutting of the donor vectors and/or inserted transgenes. The detailed generation of all plasmids used in this study is included in Table 3 of extended materials. SEED cassettes were isolated by BsmbI digestion and gel purification. Linear pUC-GW plasmids and purified SEED cassettes were assembled by NEB assembly following commercial protocols for 1h at 50°C. The samples were then dialyzed and transformed into self-prepared Top10 electrocompetent bacteria. Correct plasmids were verified by digestion and Sanger sequencing.

The 30bp-long homology arm donors were generated by PCR of the pSEED-sfGFP cassette using primers 59 and 60 and subsequent cloning into pBluescript (see Table 2 of extended materials for primer information and Supplementary figure 2B for the cloning outline). Primers included the gRNA targets for *in vivo* linearization and restriction sites SacI and EcoRI.

### Golden Gate assembly of donor SEED vectors

Golden Gate donor templates are generated as depicted in Supplementary Figure 2A. Briefly, pUC-GW plasmid containing the gRNA targets, oriented with its PAM sequence towards the homology arms, the homology arms, and cassette with two BsmBI restriction sites (sequence of the cassette: 5’*GGCTCTGAGACGCGTCTCTCCGGC3’)* was synthesized (Supplementary Figure 2A). 75ng of both the synthesized pUC-GW plasmid and the selected SEED cassette are then mixed together with one microliter of NEB Golden Gate Enzyme Mix (BsmBI-v2) in the presence of 1X T4 ligase buffer. Well mixed samples are then incubated in a thermocycler at the following temperatures: 42°C, 1 min → 16°C,1 min) x 30 → 60°C, 5 min. The samples were then dialyzed and transformed into self-prepared Top10 electrocompetent bacteria. Correct plasmids were verified by digestion and Sanger sequencing.

### Generation of gRNA expressing vectors

gRNAs were cloned into pCFD5 (Addgene 73914) by BbsI digestion and Gibson Assembly as previously described (Port & Bullock, 2016). gRNA targets can be found in Table 1 of extended materials. For gRNAs targeting *ptc* and *sqh*, transgenics were generated using the attP40 landing site (BL 25709).

### Generation of harvester stocks

pHarvester was generated using pCFD5-gRNA#1&2 and pnos-Cas9-nos plasmids (Addgene no: 62208). Briefly, the attB site in pnos-Cas9-nos was reversed by SpeI digestion and re-ligation. Subsequently, the attB, gipsy insulator, U6 promoter and gRNA array from pCFD5-gRNA#1&2 were amplified with the primers P5 and P6. Gibson Assembly was used to introduce the amplified fragment into pnos-Cas9-nos vector digested with ApaI and NheI, thus removing the mini-white marker. Harvester stocks were generated via RMCE as described previously (Sun, Johnson, Zeidler, & Bateman, 2012) using the TM3, attP.w[+].attP, Sb, Ser stock (BL 38451).

### Embryo mounting, live time-laps of dorsal closure embryos and image processing

Embryos were dechorionated in 50% bleach (sodium hypochlorite, stock solution 13% w/v technical grade; AppliChem GmbH) and washed in water. Dorsal closure stage embryos were selected manually and mounted on a glass glass-bottom dish (ibidis, 35 mm dish, ibiTreat) and embryos were covered with 1xPBS. Embryonic segment migration during dorsal closure was imaged on Point Scanning Confocal Zeiss LSM880 AiryScan inverted microscope with 63x/1.40NA objective and AiryScan detector (FAST Optimized). Time-laps series of z-stacks (12.5µm, 0.5µm z-step) with 2 minutes time resolution and 1044×1044 frame size were acquired. Raw images were AiryScan processed using Zen Black software. Z-stack maximal projections were assembled in Fiji ImageJ and signal were enhanced by multiplication.

### Fly injection

Fly embryos were injected as follows: 30 min old eggs were dechorionated in 3.5% bleach solution, aligned using a stereomicroscope and adhered on a glass slide. To avoid desiccation, embryos were covered with Voltalef H10S oil. PBS diluted samples were then injected in the posterior pole using a glass needle with the help of a pressure pump and a micromanipulator. The concentrations employed were: gRNAs plasmids, 100ng/μl; donor vectors, 100ng/μl; injections into attP lines, 300 ng/μl. We used *nos*-Cas9 flies (BL 78782) for all our knock-ins.

List of stocks used in this study

*List of stocks used in this study*
|  |  |
| --- | --- |
| *ihog*<sup>sfGFP</sup> | This study |
| *ihog*<sup>sfGFP-SEED</sup> | This study |
| *boi*<sup>mCh</sup> | This study |
| *boi*<sup>mCh-SEED</sup> | This study |
| *ptc*<sup>ALFAHA</sup> | This study |
| *sqh*<sup>ALFAHA</sup> | This study |
| *shf*<sup>sfGFP</sup> | This study |
| *shf*<sup>sfGFP-SEED</sup> | This study |
| *en*<sup>sfGFP</sup> | This study |
| *en*<sup>sfGFP-SEED</sup> | This study |
| *ihog*<sup>LOXPSEEDsfGFP</sup> | This study |
| *ihog*<sup>LOXPSEEDsfGFP-SEED</sup> | This study |
| *Dr/TM3, Sb, Ser, Harvester* | This study |
| *w*<sup>/-</sup>;M{ *UAS*-sfGFP-SEED, *w*<sup>[+]</sup> }zh-86Fb | This study |
| *w*<sup>/-</sup>;M{ *UAS*-ALFA-HA-SEED, *w*<sup>[+]</sup> }zh-86Fb | This study |
| *y*<sup>[1]</sup> *v*<sup>[1]</sup>; P{ *U6*-gRNA1+2, *v*<sup>[+]</sup> }attP40 | This study |
| *w*<sup>/-</sup>;M{ *UAS*-gRNA1+2, *w*<sup>[+]</sup> }zh-36B | This study |
| *w*<sup>/-</sup>;M{ *UAS*- gRNA1+2, *w*<sup>[+]</sup> }zh-86Fb | This study |
| *y*<sup>[1]</sup> *v*<sup>[1]</sup>; P{ *U6*-sqh gRNA 9#, *v*<sup>[+]</sup> }attP40 | This study |
| *y*<sup>[1]</sup> *v*<sup>[1]</sup>; P{ *U6*-ptc gRNAx2, *v*<sup>[+]</sup> }attP40 | This study |
| *w*<sup>/-</sup>;M{ *UAS*-deGradALFA, *w*<sup>[+]</sup> }zh-86Fb | This study |
| *w*<sup>/-</sup>;M{ *UAS*-deGradHA, *w*<sup>[+]</sup> }zh-86Fb | This study |
| *w*<sup>/-</sup>;M{ *UAS*-ER-ALFAtrap, *w*<sup>[+]</sup> }zh-86Fb | This study |
| *w*<sup>/-</sup>;M{ *UAS*-ER-ALFAtrap, *w*<sup>[+]</sup> }zh-36B | This study |
| *w*<sup>/-</sup>;M{ *UAS*-ALFAWT-mCh, *w*<sup>[+]</sup> }zh-86Fb | This study |
| *w*<sup>/-</sup>;M{ *UAS*-dALFA6mut-mCh, *w*<sup>[+]</sup> }zh-86Fb | This study |
| $w^{[-]}$;M{ *UAS-dALFA5mut-mCh*, $w^{[+]}$ }zh-86Fb | This study |
| $w^{[-]}$;M{ *UAS-dALFA3mut-mCh*, $w^{[+]}$ }zh-86Fb | This study |
| $w^{[-]}$;M{ *UAS-dALFA3mut-mGL*, $w^{[+]}$ }zh-86Fb | This study |
| $w^{[-]}$;M{ *UAS-dALFA5mut-mGL*, $w^{[+]}$ }zh-86Fb | This study |
| *ap*-Gal4 (P{GawB}*ap*<sup>md544</sup>) | Gift from Dr. Manolo Calleja |
| *nub*-Gal4 (P{GawB}*nub*-AC-62) | Gift from Dr. Manolo Calleja |
| *en*-Gal4 (P{*en2.4*-GAL4}e16E) | Bloomington number: 30564 |
| 332.3-Gal4 (P{GawB}332.3) | Bloomington number: 5398 |
| *Actin5cp4.6*-IVS-5'mCherry-#2-3'mCherry | Gift from Prof. Tzumin Lee |
| *J28-dU6-3*-gRNA(Target#2) | Gift from Prof. Tzumin Lee |
| *nos*-Cas9 | Bloomington number: 78782 |
| *elav*-Cas9 (C155) | Bloomington number: 92755 |
| *pdm2*-Gal4 | Bloomington number: 49828 |
| *UAS*-mCherry-NLS | Bloomington number: 38425 |

## Acknowledgements

Acknowledgements and founding sources

We thank Tzumin Lee, Jorge García Marques, Manolo Calleja and the Bloomington stock center for providing fly stocks and sequences used in this study, Martin Müller and Judith Cassis for their help and advice generating the *en* knock-in and Constance Cepko and Jonathan Tang for comments on the preprint. Thanks to Minkyoung Lee for her critical comments on the manuscript. The work in the laboratory of M.A. was supported by grants from the Swiss National Science Foundation (310030_192659/1) and by funds from the Kanton Basel-Stadt and Basel-Land. G.A., M.B. and S.S. were supported by ‘Fellowships for Excellence’ from the International PhD Program in Molecular Life Sciences of the Biozentrum, University of Basel. The work at I.G. laboratory was supported Grant PID2020-114533GB-C21 to IG and PRE2018-085510 to C.J-J. from the Spanish Ministry of Science, Innovation and Universities.

## Supplementary figure legends

**Supplementary figure 1. SEED reagents and validation**

**A.** Harvester stock was generated via the insertion of both *nos*-Cas9 and #1&2 into a balancer chromosome via RMCE (Sun et al., 2012). Genotype of the generated stocks is indicated. **B.** Example of 3xP3-dsRED inserted in one of the knock-ins generated in this study, in the left panel, example of larval expression of 3xP3-dsRED, strongly expressed in the mid-gut and brain. In the right panel, adult head showing strong fluorescence in the eye. **C.** Crossing scheme for the experiment of panel D. **D.** Validation of SEED cassettes. UAS-SEED cassettes were combined with harvester chromosomes and then crossed with *Act5c*-Gal4. SEED-sfGFP was scored via GFP fluorescence (n=172), ALFAHA was confirmed by PCR (n=28). In green, animals with correctly rearranged locus, in red, ds-RED positive animals, in grey, animals with not properly rearranged locus. The first bar represents the non ‘harvested’ animals, either for the sfGFP or ALFA-HA seed cassette. **E.** Crossing scheme for the generation of knock-ins using SEED/Harvest. Most often, *nosCas9* embryos are injected with gRNA-expressing vectors and SEED donors. The F0 generation is then crossed with *y,w* flies. Screening is usually done in larval stages of the F1 generation. After selection, 3xP3-expressing candidates can be crossed with balancers, to maintain the SEED cassette lines, or crossed with Harvester Balancer stocks. Cas9 is only expressed in the germ cells, making screening of dsRED/harvester candidates possible in the F2. Finally, the F2 is crossed with Balancer flies to establish the knock-in stocks. After this last step, single crosses must be established to avoid mixing of different alleles. **F.** Schematic representation of the SEED reagents. 3xP3-dsRED is flanked by the target sequences of gRNAs 1# and 2# repeated three times. The basic pSEED plasmid contains 2 MCS to facilitate downstream cloning. Asterisk: pSEED-mCherry and pSEED-mScarlet are marked with 3xP3-GFP to avoid SSA rearrangements between dsRED and the other two red fluorescent proteins.

**Supplementary figure 2. Different cloning strategies to simplify donor plasmid generation**

**A.** Scheme of the synthesis strategy. 1) Synthesis of the pUCGW vector. 185bp homology arms, flanked by targets for the gRNA were used to generate the genomic DSB (PAM orientation is highlighted in purple). In between, the right and left homology arms there are two BsmBI targets oriented tail to tail. If desired, longer linkers or other features can be added in this synthesized vector. 2) The selected SEED cassette can be inserted in a one-step Golden Gate Assembly via its BsmBI sites (see materials and methods). **B.** Addition of homology arms for MHEJ insertion of fragments with ultra-short homology. 1) SEED cassettes are amplified using forward and reverse primers that include 30bp homology arms, targets for the gRNA generation the genomic DSB and a restriction site to subclone it into pBlueScript or similar short vectors. 2) Restriction-ligation cloning into the vector. **C-D.** Details of the knock-in strategy used to generate *ptc*^ALFA:HA^ and *sqh*^ALFA:HA^. Both genes were targeted via two gRNAs provided genetically. Homology arms were designed to insert the SEED-ALFA:HA cassette before the STOP codon. **E-H.** Details of the knock-in strategy used to generate *ihog^sfGFP^*, *boi^mCh^, shf^sfGFP^* and *en^sfGFP^*. All genes are targeted using SEED cassettes bearing 185bp-long homology arms. sfGFP-SEED and mCherry-SEED cassettes are integrated in *ihog* (**E**) and *boi* (**F**), respectively, before the STOP codon. The sfGFP-SEED cassette is inserted in *shf* in the second exon (**G**) and in *en* after the first methionine (**H**).

**Supplementary figure 3. Generation of point mutations using SEED**

**A.** Scheme of the *sd* locus and the sequencing of the targeted region in exon 10. gRNAs used to trigger DSB are underlined in black (PAM is highlighted in purple). **B.** pSEED donor plasmid used as a template. The mutation is introduced in the homology arms (purple lines). Like other SEED cassettes, there is a repeat at both sides of the marker. In this case, the repeat is part of the endogenous sequence that contains the mutation. Upon HDR, the 3xP3 marker is inserted, flanked by repeated sequences. **B’.** SSA-mediated harvest leads to the assembly of the repeats, resulting in a locus modified only in the desired nucleotides. **C.** Sanger sequence of the resulting locus, showing the desired mutation (highlighted in red) and the two silent mutations introduced to avoid re-cutting by the gRNAs.

**Supplementary figure 4. Tissue-specific activation of SEED cassettes in the brain and updated SEED vectors**

**A.** Confocal image of the larval VNC of *ihog*^SEED-sfGFP^ flies combined with *vGlut*-Cas9 and ubiquitously expressed gRNA#2. Ihog labels small structures that resemble somas at the lateral regions. **B.** Confocal image of the larval VNC of *ihog*^SEED-sfGFP^ flies combined with *elav*-Cas9 and ubiquitously expressed gRNA#2. Similar to *vGlut*-Cas9, somas can be labelled. **B’.** Detail magnified view of some of the somas labelled with Ihog using *elav*-Cas9. **C.** Wing disc displaying clones of Ihog-sfGFP expression. From the same genotype as in B and B’. **D.** Scheme of the pSEED-LoxP-FP vectors. Two loxPs are inserted flanking the 3xP3-FP marker. **E.** Upon Cre activation, the LoxP sites recombine, removing the 3xP3 marker. The cassette remains sensitive to the activation via both gRNAs #1 and #2. **F.** Percentage of founder 3xP3-dsRED animals using pSEED-Ihog-sfGFP and pSEED-ihog-sfGFP-LoxP. pSEED-Ihog-sfGFP: 43% (n=28), pSEED-ihog-sfGFP-LoxP 42% (n=24). Scale bars: 100μm.

**Supplementary figure 5. Hh signaling changes upon *shf^S^*^EED-sfGFP^ rescue**

**A**. anti-Ptc immunostaining (green) of *shf^S^*^EED-sfGFP^ wing discs. Ptc appears localized in the anterior compartment, in a row of cells in contact with the posterior compartment. Low levels are detected through the rest of the anterior compartment. **A’.** anti-Ci^155^ immunostaining (red) of *shf^S^*^EED-sfGFP^ wing discs, staining is, as Ptc, constrained to the first cell row of the anterior compartment. **B**. anti-Ptc immunostaining (green) of *shf^s^*^fGFP^ wing discs. Ptc is detected at high levels in the first 4 cell rows of the anterior compartment in contact with the posterior compartment. Levels through the rest of the anterior compartment are low but higher than in the panel A. **B’.** anti-Ci^155^ immunostaining (red) of *shf^s^*^fGFP^ wing discs, Ci extends in a broad stripe along the anterior compartment. In turn, high Hh levels close to the compartment border cause its repression in the cell rows most proximal to the posterior compartment (arrowhead). **C**. anti-Ptc immunostaining (green) in wing discs of *shf^S^*^EED-sfGFP^ animals rescued in the *nub*-Gal4 domain. The affected domain is outlined with red dots. Ptc localizes at high levels in the first 4 cell rows of cells of the anterior compartment in contact with the posterior compartment. Levels through the rest of the anterior compartment are low but higher than in the panel B. **C’.** anti-Ci^155^ immunostaining (red) of *shf^s^*^fGFP^ wing discs, Ci extends in a broad stripe along the anterior compartment. In turn, high Hh levels close to the compartment border cause its repression in the cell rows most proximal to the posterior compartment (arrowhead). **D**. anti-Ptc immunostaining (green) in wing discs of *shf^S^*^EED-sfGFP^ animals rescued in the *ap*-Gal4 domain. Affected domain is outlined with red dots. Ptc levels are higher than the panel A, but never as high as in B or C. Levels in the dorsal compartment are higher than those in the ventral compartment. Levels through the rest of the anterior compartment are low but higher than in the panel B. **D’.** anti-Ci^155^ immunostaining (red) of *shf^s^*^fGFP^ wing discs, Ci extends in a broad stripe along the anterior compartment. Levels are generally lower than those of the panels B and C, but the Hh-signaling range is clearly rescued compared to panel A. In the ventral compartment, far from the dorso-ventral compartment border, the stripe appears to be less broad than in dorsal compartment. High Hh levels close to the compartment border still cause its repression in the cell rows most proximal to the posterior compartment (arrowhead). **E.** Detailed magnification of Ptc immunostaining depicted in D. Note the higher range of the dorsal signal vs the ventral signal **F-I.** Image reconstruction of an image stack of the heads of the genotypes depicted in A-D. Notice the reduced size of the eyes in images F, H and I compared to G (*shf^s^*^fGFP^). Quantified in Figure 4I. **F’-I’.** Close-up image reconstruction of an stack of images of the eyes depicted in F-I. Notice the ordered pattern of G’ compared to the often-fused ommatidia (arrowheads) displayed by F’, H’ and I’. Scale bars: A-D: 100μm, E: 25μm.

**Supplementary figure 6. Simultaneous visualization and ER-trapping of double-tagged knock-ins**

**A.** Scheme of the ALFA-Trap^ER^. CD8 (black rectangular box) is fused to mCherry (red circle) at its luminal/extracellular region and to ALFANb (yellow oval) at its cytoplasmic tail. In addition, KKXX sequence was added in CD8’s C-terminus to mediate retention in the ER. If co-expressed with ALFA-tagged transmembrane peptides (in this case, the 12-pass transmembrane protein Ptc) ALFA-Trap^ER^ can impede its secretion. **B.** Wing discs of *ptc*^ALFA:HA^ flies in absence (first column) or presence (second column) of the ALFA-Trap^ER^. The mCherry signal is observed in the wing pouch upon ALFA-Trap^ER^ expression (in red). anti-HA immunostaining, reveals a stripe of cells across the antero-posterior boundary expressing Ptc (first column). Increased levels of Ptc:ALFA:HA in the wing disc are apparent upon expression of ALFA-Trap^ER^. Scale bars: 100μm.

