## Supplementary Figure for "*In vivo* seamless genetic engineering via CRISPR-triggered single-strand annealing"

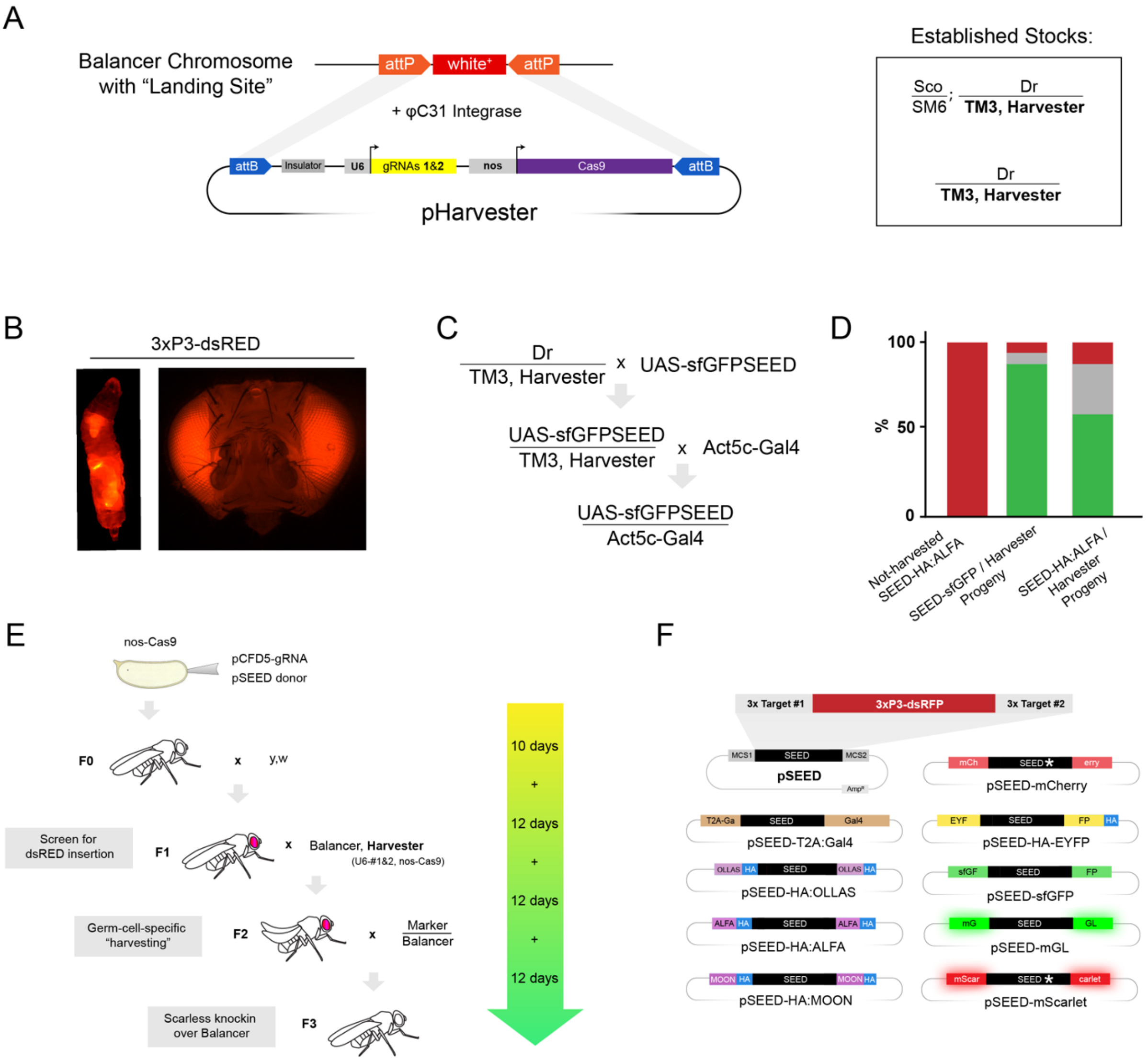


**Supplementary figure 1. SEED reagents and validation**

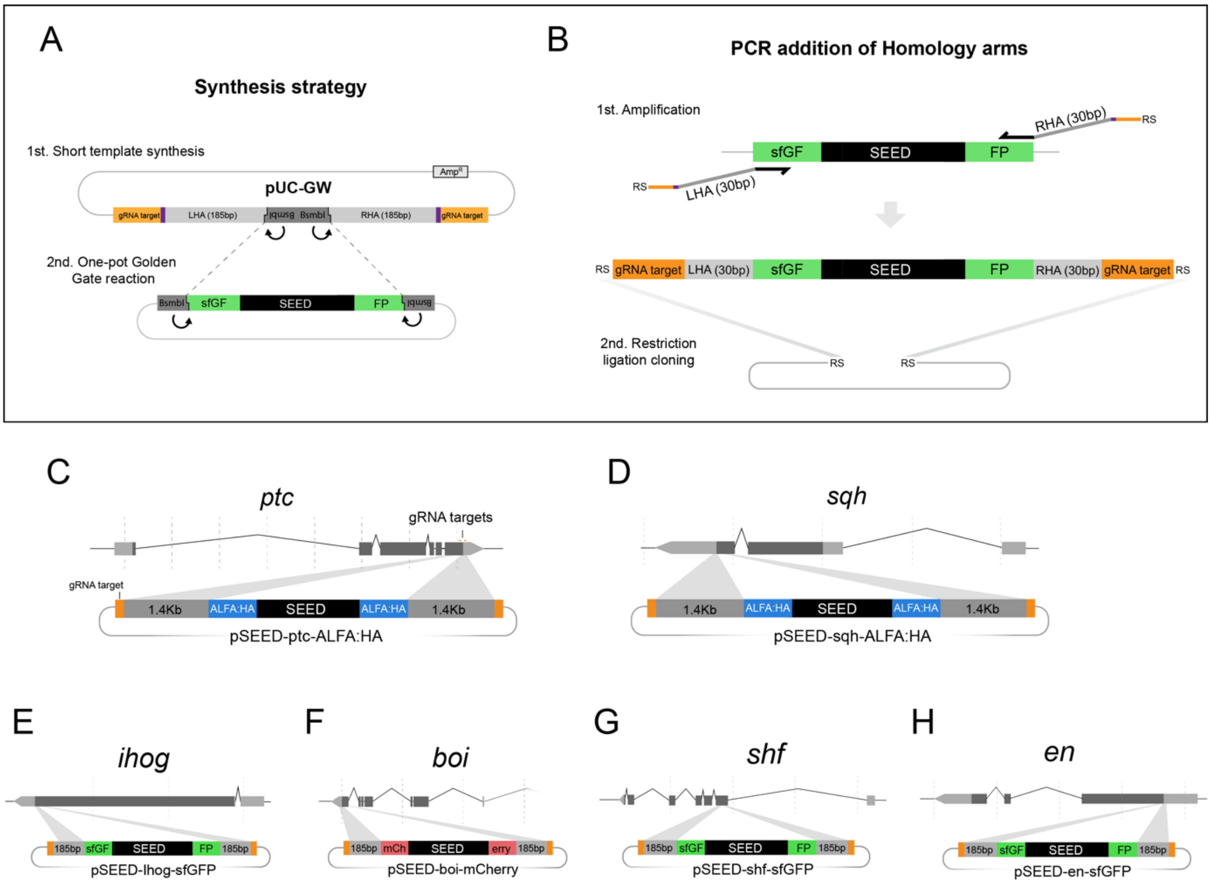


**Supplementary figure 2. Different cloning strategies to simplify donor plasmid generation.**

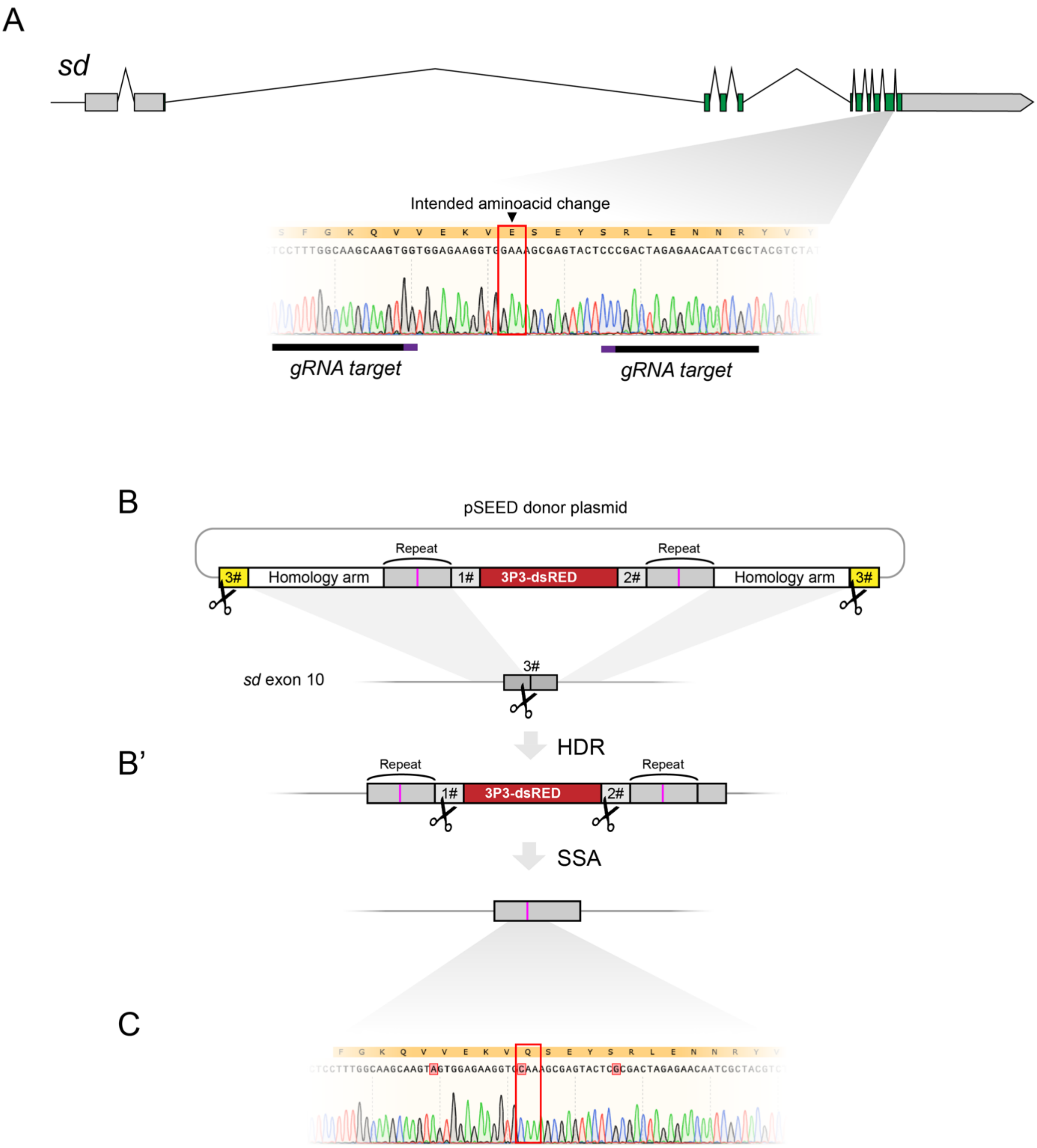


**Supplementary figure 3. Generation of point mutations using SEED**

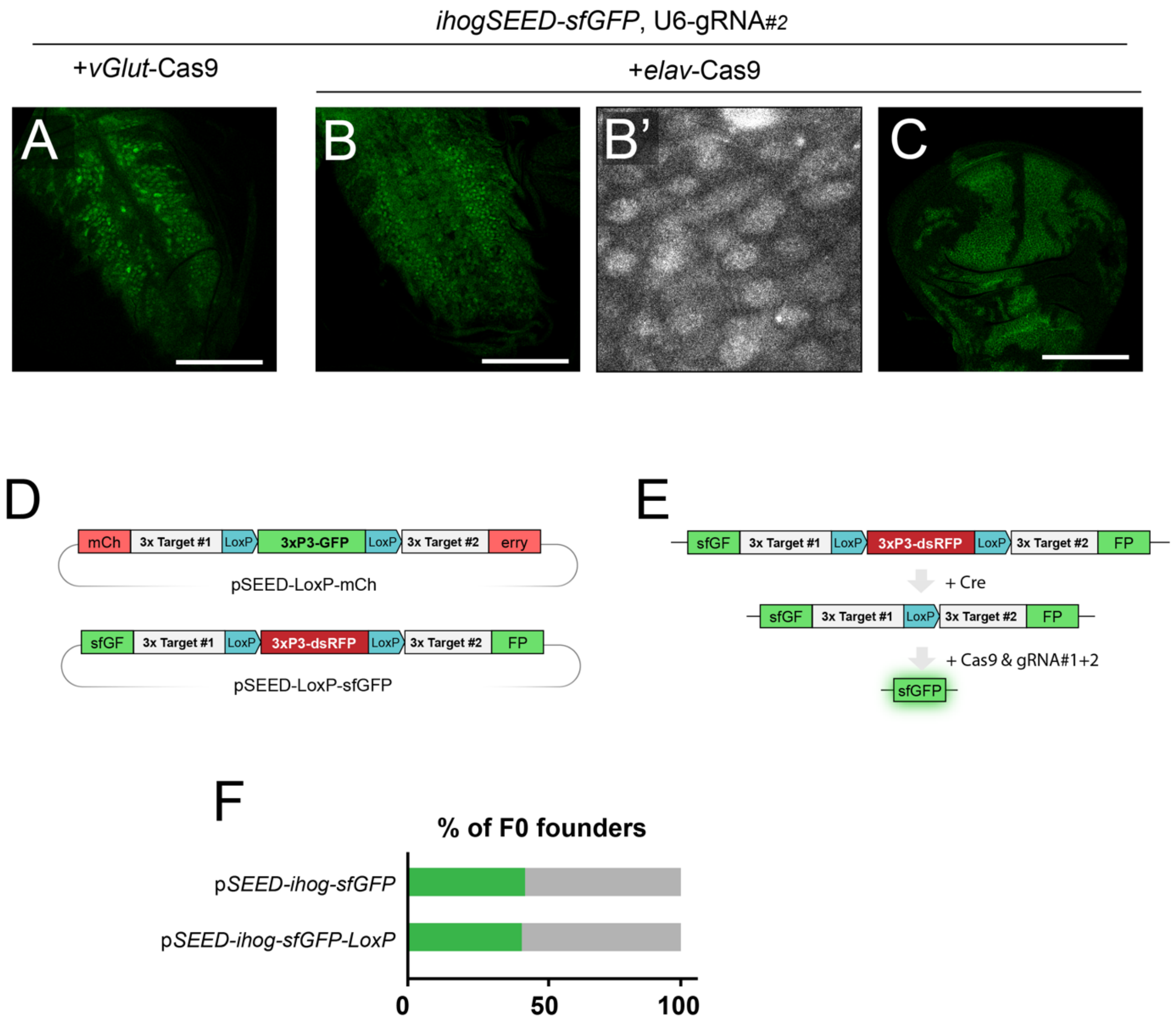


**Supplementary figure 4. Tissue-specific activation of SEED cassettes in the brain and updated SEED vectors**

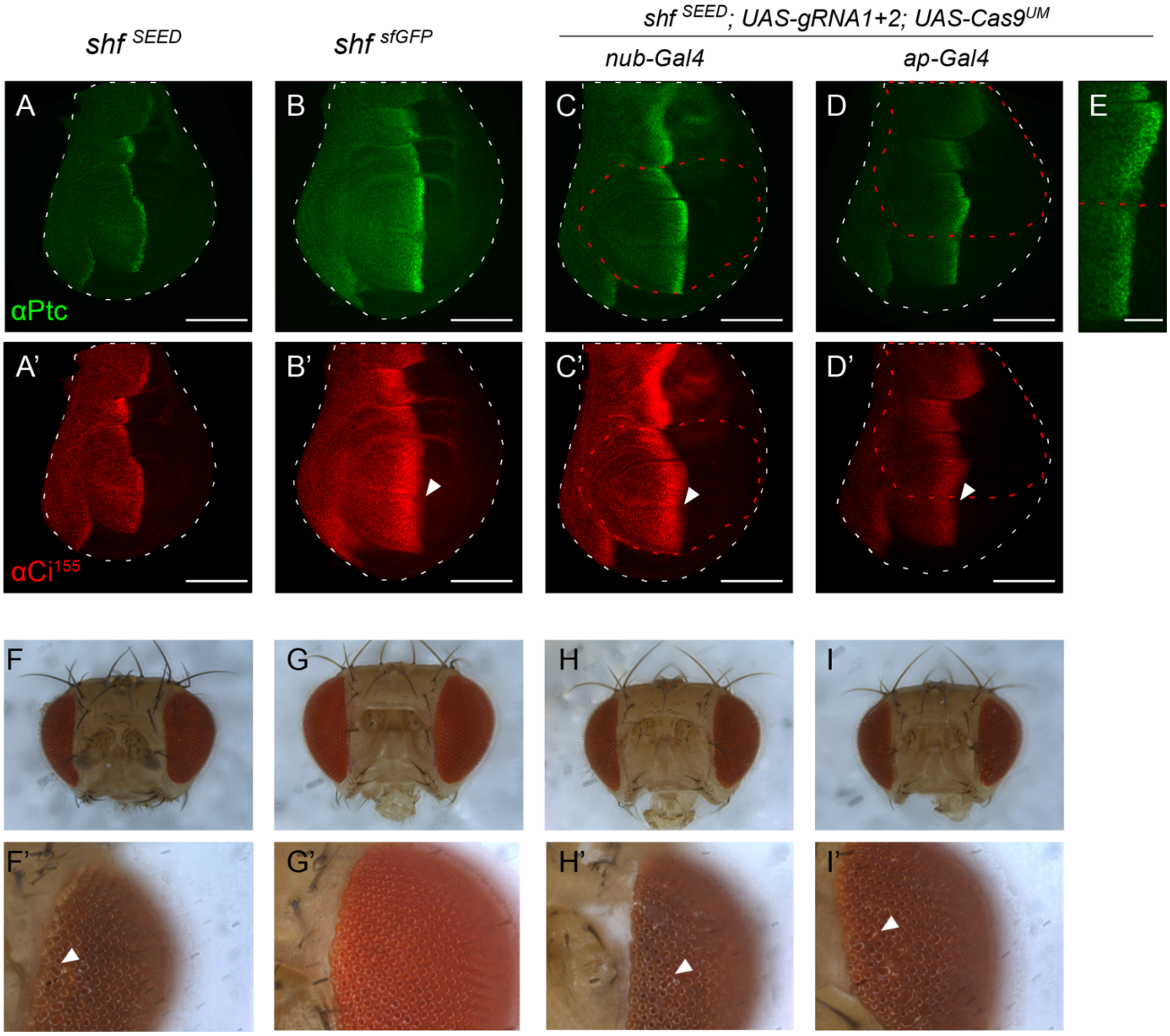


**Supplementary figure 5. Hh signaling changes and phenotypes upon *shf*$^{\mathrm{SEED\text{-}sfGFP}}$ rescue**

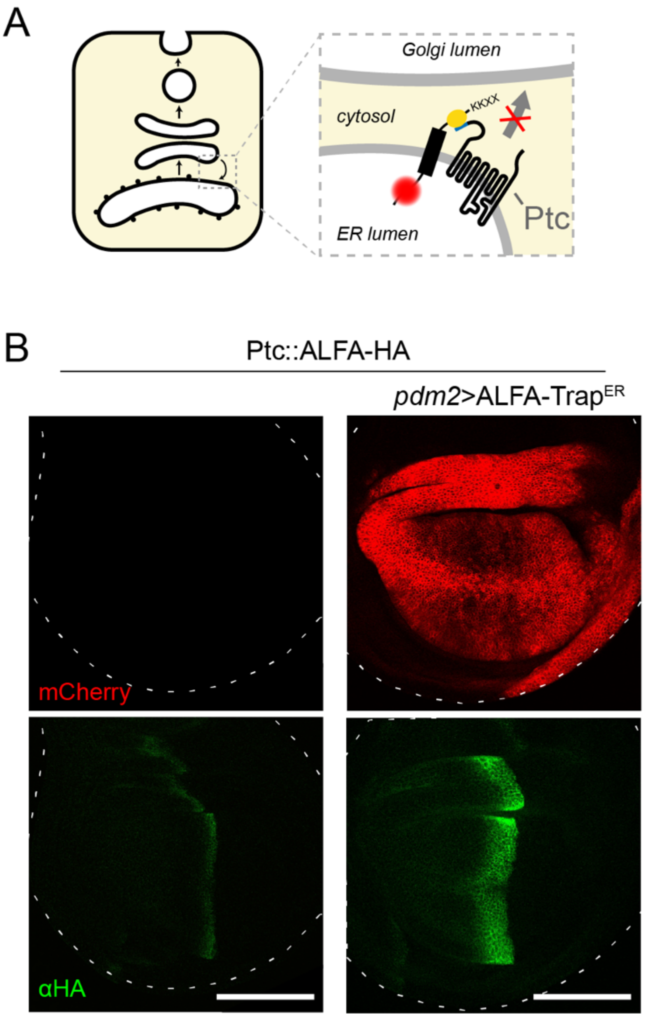


**Figure 6. Simultaneous degradation and visualization of double-tagged knock-ins**
