## Extended Materials for "*In vivo* seamless genetic engineering via CRISPR-triggered single-strand annealing"

**Extended Materials. Aguilar et al. 2022.**

Synthesized fragments:

#### Shf SEED

GAAGTTGTCCAGGCCATCGCTGGAAACTATGACACATCAGGGCATCGGCTGCCTGGTCAAGTGGTTATATCTGGTGCTGATCGTGCACACGCTCCTCTGCATCGGCCAGCTCGAGTGTCGCCAGCAGCACCACAATCGGAATAACAATAATAATAACAGAAGAGCGGACTCGTCCAGCTCAGAGGAGGGCCACGGCAACACCAGCGATGGCAGTGGCAGTATGGTGTCCAAGGGCGAGGAGCTGTGGTCTTCGAAGACACCTGGGCATGGATGAGCTGTACAAGTCCGGCGGCAGTGGCCTGGACAACTTCGCAGATCAGGACGCCAGCTTCGTTGGCCATGGACACCAGCCGCGTCGGGGTCAACGAAAGAAACAGCAGGGAGGTGGAGGAGGAGGCAGCGGCGGCGGAAACGGAGGAGGCGGTGGAAGCCGGCACAATCGCAACGAGGAGAGCGGCATCTCGTTGTGGATCAATGAGCACCAGCGATGGCCTGGACAACTTC

#### Boi SEED-mCh

TGAACAGTATTGAGGTGTGATGGCGCACCACCAGCATCTGCACCACCACCCCGGCCATCCGGGTGGCCTGGTGGGCATACCCATCGTGCCAGGGAGCCCTAGGGTCCAGAGATCGCCGATGCCGAGCAGGGCGATGATCAAGAGGACGCGCCTGGGCAGCCACACGGACAACATATCGTCGGGCAGTCTGAACAGTATTGAGGTGGGTGGAGGCTCTATGGTGAGCAAGGGCGAGGAGGATTGGTCTTCGAAGACACGGCGGCATGGACGAGCTGTACAAGTCCGGCTGATGGCTCGCAGAGGAGGCAACTAACTAGTGGACTAGTGGGCGGAGCAGCACAACGAGGAGAAATATACCTAAGCGGCAAGCGAAAGTGGTTCCCCAAAAACAGACAGTTTCAAGTAGACTTAAGGATCCGAAACTAGATGTGTATTATTGAGATAATCGTAGGACAGGTCGGCAGAGCAGCTCGAACCATCACACCTCAATACTGTTCA

#### Ihog-SEED-sfGFP

TAGCATGGTACATCGGGCGTAGGCGTCGCTCACAACGCACGCTGGAACGTGCTGGCGGCAGCAATGGCTCCAACAACGGCAACAACAACAATCTAAATCAATCCGCAGAGGCGGGTTCCATCGAAAATCCCGGCAAGCCGGGTCGTGTACTCATGAAACGCCCCCGTCTGTCCAGTCGCTCCGAGAACCTCTCCTCCGGCAGtCTCAACAGCGTTGGCGTGGGTGGAGGCAGTGGCAGTATGGTGTCCAAGGGCGAGGAGCTGTGGTCTTCGAAGACACCTGGGCATGGATGAGCTGTACAAGTCCGGCTAATCGCAATCTCGTGATCTTTAGTCTGTGCATCACTACGCCCGATGTACCATGCTATTCCTCTACCTCTGTTTTATCTACCACTTTGAATGACAAAACCTTTGCTGTTTGTACTTAAATAGGTACTTAAATAGACATAACTAGATTTATGTATGCATGATCGTTTAAGAACATAGCATTATGTCCATTCTACGCTAAcTCTTCCATTTTTCGCGATAAGATCAAAA CCTACGCCCGATGTACCATGCTA

#### en-SEED-sfGFP

CAAGTCGAAACCAATGGCCCTGGCAGACGTGCGTGTCGGAACAACAGTTGCAAATCAAACACGAAAGCATAAGCCAAACAAAAAACACCAAACAGAGAAGAGAATCAGAAGATTCTGAGCAATCAGAAGAATCAGTGGCTCAGTGTCAAGTGACCCAGTGACAAGTGTCTTAAGCGAGTTGCGATTTAGCACCAAGTCGAAACCAATGGGTGGAGGCAGTGGCAGTATGGTGTCCAAGGGCGAGGAGCTGTGGTCTTCGAAGACACCTGGGCATGGATGAGCTGTACAAGTCCGGCGCCCTGGAGGATCGCTGCAGTCCACAGTCAGCGCCCAGCCCCATTACCCTACAAATGCAGCATCTTCACCACCAGCAACAGCAGCAGCAGCAACAGCAGCAGCAAATGCAGCACCTCCACCAGCTGCAGCAACTGCAGCAGTTGCACCAACAGCAACTGGCCGCCGGTGTCTTCCACCATCCGGCCCAGGGCCATTGGTTTCGACTTG

gRNA targets

185bp Homology Arms

Linker sequences

Homology for Gibson assembly with the SEED cassette chosen in each case

#### ALFANbWT

CAAAATGAGCGGAGAGGTGCAGCTGCAGGAGAGCGGCGGCGGCCTGGTGCAGCCCGGCGGCAGCCTGCGCCTGAGCTGCACCGCCAGCGGCGTGACCATCAGCGCCCTGAACGCCATGGCCATGGGCTGGTATCGCCAGGCCCCCGGCGAGCGCCGCGTGATGGTGGCCGCCGTGAGCGAGCGCGGCAACGCCATGTACCGCGAGAGCGTGCAGGGCCGCTTCACCGTGACCCGCGATTTCACCAACAAGATGGTGAGCCTGCAGATGGATAACCTGAAGCCCGAGGATACCGCCGTGTACTACTGCCACGTGCTGGAGGATCGCGTGGATAGCTTCCACGATTACTGGGGCCAGGGCACCCAGGTGACCGTGAGCAGCGGAGGCGGTAGCGGTAAGCCTATCCCTAACCCTCTCCTCGGTCTCGATTCTACGGGCGGTGGATCC

#### ALFANb3mut

CAAAATGAGCGGAGAGGTGCAGCTGCAGGAGAGCGGCGGCGGCCTGGTGCAGCCCGGCGGCAGCCTGCGCCTGAGCTGCACCGCCAGCGGCGTGACCATCAGCGCCCTGAACGCCATGGCCATGGGCTGGTATCGCCAGGCCCCCGGCGAGCGCCGCGTGATGGTGGCCGCCGTGAGCGAGCGCGGCAACGCCATGTACCGCGAGAGCGTGCAGGGCCGCTTCACCGTGCGCCGCGATTTCACCAACAAGATGGTGAGCCTGCAGATGGATAACCTGAAGCCCGAGGATACCGCCGTGTACTACTACCACGTGCTGGAGGATCGCGTGGATAGCTTCCACGATTACTGGGGCCAGGGCACCCAGGTGACCGTGAGCTTCGGAGGCGGTAGCGGTAAGCCTATCCCTAACCCTCTCCTCGGTCTCGATTCTACGGGCGGTGGATCC

#### ALFANb5mut

CAAAATGAGCGGAGAGGTGCAGCTGCAGGAGAGCGGCGGCGGCCTGGTGCAGCCCGGCGGCAGCCTGCGCCTGAGCTGCGTTGCCAGCGGCGTGACCATCAGCGCCCTGAACGCCATGGCCATGGGCTGGTATCGCCAGGCCCCCGGCGAGCGCCGCGTGATGGTGGCCGCCGTGAGCGAGCGCGGCAACGCCATGTACCGCGAGAGCGTGCAGGGCCGCTTCACCGTGCGCCGCGATTTCACCAACAAGATGGTGAGCCTGCAGATGGATAACCTGAAGCCCGAGGATACCGCCGTGTACTACTACCACGTGCTGGAGGATCGCGTGGATAGCTTCCACGATTACTGGGGCCACGGCACCCAGGTGACCGTGAGCTTCGGAGGCGGTAGCGGTAAGCCTATCCCTAACCCTCTCCTCGGTCTCGATTCTACGGGCGGTGGATCC

#### ALFANb6mut

CAAAATGAGCGGAGAGGTGCAGCTGCAGGAGAGCGGCGGCGGCCTGGTGCAGCCCGGCGGCAGCCTGCGCCTGAGCTGCGTTGCCAGCGGCGTGACCATCAGCGCCCTGAACGCCATGGCCATGGGCTGGTATCGCCAGGCCCCCGGCGAGCGCCGCGTGATGGTGGCCGCCGTGAGCGAGCGCGGCAACGCCATGTACGTGGAGAGCGTGCAGGGCCGCTTCACCGTGCGCCGCGATTTCACCAACAAGATGGTGAGCCTGCAGATGGATAACCTGAAGCCCGAGGATACCGCCGTGTACTACTACCACGTGCTGGAGGATCGCGTGGATAGCTTCCACGATTACTGGGGCCACGGCACCCAGGTGACCGTGAGCTTCGGAGGCGGTAGCGGTAAGCCTATCCCTAACCCTCTCCTCGGTCTCGATTCTACGGGCGGTGGATCC

5' Alfa HA (250bp)

CAATTTCACACAGGAAACAGCTATGACCATGATTACGCCAGAGCTCACGCGTACCGGTCTCGAGGAATTCGCCGGCGGAGGATCTGGACCATCACGTTTGGAAGAGGAACTGAGACGCCGCTTAACTGAACCTGGGGCAGGATACCCATACGATGTTCCAGATTACGCTGGATCCGGCCCTAGGCCGCGTTGTTATGATCGTACTACGCTAGCGTAGTACGATCATAACAACGCGGGTAGTACGATCATA

#### 3' Alfa HA (245bp)

ACTGTATGGATGTACCCGCTGGTACTGTATGGATGTACGCTAGCGTACATCCATACAGTACCAGCGGACTAGTGGCGGAGGATCTGGACCATCACGTTTGGAAGAGGAACTGAGACGCCGCTTAACTGAACCTGGGGCAGGATACCCATACGATGTTCCAGATTACGCTGGATCCGGCGCCGCATGCCTTAAGAGATCTGGTACCTGGTGCACTCTCAGTACAATCTGCTCTGATGCCGCATAGT

Table 1. GuideRNAs used in the study.

| **Targeted locus** | **gRNA1** | **gRNA2** |
| --- | --- | --- |
| *ptc* | GCTGGCCATGCCCGGCAGGG | TGTAAAATCGATTTGTCCAG |
| *ihog* | ATTACACGCCAACGCTGTTG | TAGCATGGTACATCGGGCGT |
| *boi* | TGAACAGTATTGAGGTGTGA |  |
| *sqh* | CATCCTTAAGCACGGTGCCA |  |
| *shf* | GAAGTTGTCCAGGCCATCGC |  |
|  | CAAGTCGAAACCAATGGCCC |  |
| *gRNA1#* | GTAGTACGATCATAACAACG |  |
| *gRNA2#* | GTACATCCATACAGTACCAG |  |
| *sd* | TTGCTCCTTTGGCAAGCAAG | ACGTAGCGATTGTTCTCCAG |

Table 2. List of primers used in this study.

| **Primer number (#)** | **Primer name** | **PRIMER SEQUENCE** |
| --- | --- | --- |
| **P1** | For pSEED sfGFP | GGTGGAGGCAGTATGGTGTCCAAGGG |
| **P2** | Rev pSEED sfGFP | GCCGGACTTGTACAGCTCATC |
| **P3** | Rev pSEED mCherry | GGTGGAGGCTCTATGGTGAGCAAGGG |
| **P4** | Rev pSEED mCherry | GCCGGACTTGTACAGCTCG |
| **P5** | Gibson 1+2 For | CTATAGGGCGAATTGGGTACGTACCGGGCCCCGACGATGTAGGTCACGG |
| **P6** | Gibson 1+2 Rev | GGTTAAAACGGTCGAAGCTTGGATCCGGAACAAAAGCTGGAGCTCCTG |
| **P7** | pLOTfor | GAACTCTGAATAGGGAATTGGG |
| **P8** | Reverse Cd8 | CTGTGGTAGCAGATGAGAGTG |
| **P9** | Alpha Nb For Gibson | CTGCTGTCCTTGATCATCACTCTCATCTGCTACCACAGCGGAAGCGAGGTGCAGCTGCAGG |
| **P10** | AlfaNb.KKXX.STOP Gibson | ACACCACAGAAGTAAGGTTCCTTCACAAAGATCCTCTAGATTATAGGTGCTTCTTTCCGCTGCCTCCGCTGCTCACGGTCACC |
| **P11** | F guide1 | GCGGCCCGGGTTCGATTCCCGGCCGATGCAGTAGTACGATCATAACAACGGTTTTAGAGCTAGAAATAGCAAG |
| **P12** | Reverse guide2 | ATTTTAACTTGCTATTTCTAGCTCTAAAACCTGGTACTGTATGGATGTACTGCACCAGCCGGGAATCGAACCC |
| **P13** | For Ihog gRNA | GCGGCCCGGGTTCGATTCCCGGCCGATGCAATTACACGCCAACGCTGTTGGTTTTAGAGCTAGAAATAGCAAG |
| **P14** | Rev Ihog gRNA | ATTTTAACTTGCTATTTCTAGCTCTAAAACACGCCCGATGTACCATGCTATGCACCAGCCGGGAATCGAACCC |
| **P15** | F Boi gRNA | TGCATGAACAGTATTGAGGTGTGA |
| **P16** | R Boi gRNA | AAACTCACACCTCAATACTGTTCA |
| **P17** | F Shf | TGCAGAAGTTGTCCAGGCCATCGC |
| **P18** | R Shf | AAACGAAACTATGACACATCAGGG |
| **P19** | ptcF | GCGGCCCGGGTTCGATTCCCGGCCGATGCAGCTGGCCATGCCCGGCAGGGGTTTTAGAGCTAGAAATAGCAAG |
| **P20** | ptcR | ATTTTAACTTGCTATTTCTAGCTCTAAAACCTGGACAAATCGATTTTACATGCACCAGCCGGGAATCGAACCC |
| **P21** | F ApaI | TAGGGCCCCGACGATGTAGGTCACGG |
| **P22** | R Nhe1 | TAGCTAGCGAACAAAAGCTGGAGCTCCTG |
| **P23** | For sfGFP 1 EcoRI | AGGAATTCGCCGGCAGTATGGTGTCCAAGGGCG |
| **P24** | Rev sfGFP AvrII | AGCCTAGGGCGGATCTTGAAGTTGGC |
| **P25** | Rev sfGFP Sph1 | AGGCATGCGGCGCCGGACTTGTACAGCTCATCCATGCC |
| **P26** | For sfGFP spe1 | CAACTAGTCCAACGGCAAGCTGAC |
| **P27** | Cassette reverse SalI | ACGTCGACGTACATCCATACAGTACCAGCGGGTACATCCATACAGTACCAGCGGTAAGATACATTGATGAGTTTGGACAAACCAC |
| **P28** | Cassette forward Nhe1 | ACGCTAGCGTAGTACGATCATAACAACGCGGGTAGTACGATCATAACAACGCGGGGATCTAATTCAATTAGAGACTAATTCAATTAGAGC |
| **P29** | mCherryPartIF | GAGGAATTCGCCGGCTCTATGGTGAGCAAGGGCGA |
| **P30** | mCherryPartIRnew | GCGGCCTAGGGCCGGAGCCGCCGTCCTTCA |
| **P31** | mCherryPartIIF | CGGACTAGTGGCTCTCGAGGGCACCCAGAC |
| **P32** | mCherryPartIIR | AAGGCATGCGGCGCCGGACTTGTACAGCTCGTCCATGC |
| **P33** | ollasHATagR2 | CGGCGCCGGATGCATAGTCCGGGACGTCATAGGGATATTTACCCATCAGGCGGGGTCCCAGCTCGTTCGCGAAGCCGCTAGAGCCA |
| **P34** | OllasHAF2 | CTAGTGGCTCTAGCGGCTTCGCGAACGAGCTGGGACCCCGCCTGATGGGTAAATATCCCTATGACGTCCCGGACTATGCATCCGGCGCCGCATG |
| **P35** | ollasHAdoubleF | AATTCGCCGGCTCTAGCGGCTTCGCGAACGAGCTGGGACCCCGCCTGATGGGTAAATATCCCTATGACGTCCCGGACTATGCATCCGGCC |
| **P36** | ollasHATagR | CTAGGGCCGGATGCATAGTCCGGGACGTCATAGGGATATTTACCCATCAGGCGGGGTCCCAGCTCGTTCGCGAAGCCGCTAGAGCCGGCG |
| **P37** | ecor1testmoontagfor | GAGGAATTCGCCGGCGGAGGATCTGGAAAGAACGAGCAGGAACTGCTGGAGC |
| **P38** | testecoN1moontagrev | GATCGTATCCTGCCCCAGGCAGAGAGGCCCATTTGTCCAGCTC |
| **P39** | spetestmoontagfor | GATCACTAGTGGCGGAGGATCTGGAAAGAACGAGCAGGAACTGCTGGAGC |
| **P40** | testecoN1moontagrev | GATCGTATCCTGCCCCAGGCAGAGAGGCCCATTTGTCCAGCTC |
| **P41** | For T2A Gal4 EcoR1 | AGGAATTCGCCGGCGAGGGCAGGGGAAGTCTTCTAACATGCGGGGACGTGGAGGAAAATCCCGGCCCCATGAAGCTACTGTCTTCTATCGAAC |
| **P42** | Reverse Gal4 AvrII | TACCTAGGCCGATGATGATGTCGCAC |
| **P43** | For Gal4 Spe1 | CTACTAGTATGAAGCTACTGTCTTCTATCGAAC |
| **P44** | Rev Gal4 Kpn1 | ATGGTACCTTACTCTTTTTTTGGGTTTGGTGGG |
| **P45** | Ptc Rev HA1 SSA | TTCCAAACGTGATGGTCCAGATCCTCCGCCACTCGTAAAGTTATAGCTGCGCA |
| **P46** | ptc For HA1 SSA | ACGCGTACCGGTCTCGAGGAATTCGCCGCTGGCCATGCCCGGCAGGGCGGCAACGAGTACGATCTTAAGATACCC |
| **P47** | For HA2 SSA | TACGATGTTCCAGATTACGCTGGATCCGGCTAGCACTAGCACTAGTTCCTGTAG |
| **P48** | Reverse Ptc 1 | CGAGCTACAATTACTAGTTTG |
| **P49** | Forward ptc seq2 | CAAACTAGTAATTGTAGCTCG |
| **P50** | Ptc Rev HA2 SSA | CCAGGTACCAGATCTCTTAAGGCATGCGGCCCGCCCTGCCGGGCATGGCCAGCCAAATCTGATCAATTGACTAAATCACC |
| **P51** | For dsRED LoxP | TACGCTAGCGTAGTACGATCATAACAACGCGGGTAGTACGATCATAACAACGCGGATAACTTCGTATAGCATACATTATACGAAGTTATGGATCTAATTCAATTAGAGACTAATTC |
| **P52** | Rev LoxP dsRed | TACGCTAGCGTACATCCATACAGTACCAGCGGGTACATCCATACAGTACCAGCGGATAACTTCGTATAATGTATGCTATACGAAGTTATTAAGATACATTGATGAGTTTGGACAAACC |
| **P53** | nbalfashxho_for | GATCCTCGAGGAGGTGCAGCTGCAGGAG |
| **P54** | nbalfasxba_rev | GATCTCTAGACTATGAGGAGACGGTGACC |
| **P55** | eyfp5phlfecor1_for | GATCGGAATTCGGAGGAGGGATGGTGAGCAAGGGC |
| **P56** | eyfp5pavr2_rev | GATCCCTAGGCCTCGATGTTGTGGCGGATCTTG |
| **P57** | eyfp3pspe_for | GATCACTAGTCCTACGGCAAGCTGACCCT |
| **P58** | haeyfp3psph1_rev | GATCGCATGCTTATCCGGATCCAGCGTAATC |
| **P59** | Rev Boi 30bp HA R mCh | TACGAATTCTGAACAGTATTGAGGTGTGATGGCTAGTTAGTTGCCTCCTCTGCGAGCCATCAGCCGGACTTGTACAGCTC |
| **P60** | For Boi 30bp HA F mCh | TACGAGCTCTGAACAGTATTGAGGTGTGATGGTCGTCGGGCAGTCTGAACAGTATTGAGGTGGGTGGAGGCTCTATGGTGAGCAAGG |
| **P61** | sqh5F | TGCACATCCTTAAGCACGGTGCCA |
| **P62** | sqh5R | AAACTGGCACCGTGCTTAAGGATG |
| **P63** | sdguide1F | TGCACGTAGCGATTGTTCTCTAGT |
| **P64** | sdguide1R | AAACACTAGAGAACAATCGCTACG |
| **P65** | sdguide2F | TGCATTGCTCCTTTGGCAAGCAAG |
| **P66** | sdguide2R | AAACCTTGCTTGCCAAAGGAGCAA |
| **P67** | sd For HA1 Age1 | CGCGTACCGGTTTGCTCCTTTGGCAAGCAAGTGGGGCAAATATTCGACAAGTTTCCGGAG |
| **P68** | sd HA1 rev AvrII | GGCTACCTAGGGGGAGCGTTGAATGCGATAGACGTAGCGATTGTTCTCTAGTCGCGAGTACTCGCTTTGCACCTTCTCCATCACTTGCTTGCCA |
| **P69** | sd For HA2 SpeI | AGCGGACTAGTAATGTCGTGCTCGTGTGCTCCACAATCGTTTGCTCCTTTGGCAAGCAAGTGATGGAGAAGGTGCAAAGCGAGTACTCGCGACTAGAG |
| **P70** | sd Rev HA2 KpnI | CACCAGGTACCCGTAGCGATTGTTCTCTAGTCGGTGTGAACTTGTGTGTTGATGATGGG |

Table 3. Plasmids generated in this study and generation procedure.

| pUASattb-ALFAHA-SEED: | Generated by subcloning the ALFAHA-SEED cassette from the pSEED-ALFAHA digesting both with EcoRI and KpnI. |
| --- | --- |
| pUASattB-sfGFP-SEED: | Generated by subcloning the sfGFP-SEED cassette from the pSEED-ALFAHA digesting both with EcoRI and KpnI. |
| pUASattB-ALFANb:mCherry | Generated by fragment synthesis of the ALFANbWT fragment and subcloning into pUAS-mCherry plasmid via EcoRI and KpnI |
| pUASattB-dALFANb(6mut):mCherry | Generated by fragment synthesis of the ALFANb6mut fragment and subcloning into pUAS-mCherry plasmid via EcoRI and KpnI |
| pUASattB-dALFANb(5mut):mCherry | Generated by fragment synthesis of the ALFANb5mut fragment and subcloning into pUAS-mCherry plasmid via EcoRI and KpnI |
| pUASattB-dALFANb(3mut):mCherry | Generated by fragment synthesis of the ALFANb3mut fragment and subcloning into pUAS-mCherry plasmid via EcoRI and KpnI |
| pLOT(LexO/UAS)-mCherry:CD8:ALFANb:KKXX | Generated by Gibson Assembly of mCherry:CD8 and ALFA Nb fragments (amplified from morphotrapint with P7-P8 primers and from pUASattB-ALFANb:mCherry with P9-P10 primers, respectively) into pLOT-morphotrapInt (Harmansa et al, 2017) digested with XhoI and XbaI |
| pUASattB-deGradALFA | The VHH4 nanobody of pUASTattB_NSlmb-vhhGFP4 (E. Caussinus et al., 2011) was substituted by ALFA nanobody from pNT-NAM01 pCMV-NbALFA- MCS plasmid (Gift by S. Frey). ALFANb was amplified using P53 and P54 and then digested for subsequent ligation using XhoI and XbaI. |
| pCFD5-gRNA#1&2: | Generated by Gibson Assembly into the BbsI-digested pCFD5 vector using the primers P11 and P12. |
| pCFD5-sqh-gRNA#5 | Generated by ligation into BbsI-digested pCFD5 vector of the annealed primers P61 and P62 |
| pCDF5-ihog-2xgRNA-CTerm: | Generated by Gibson Assembly into the BbsI-digested pCFD5 vector using the primers P13 and P14. |
| pCDF5-boi-gRNA-CTerm: | Generated by ligation into BbsI-digested pCFD5 vector of the annealed primers P15 and P16 |
| pCDF5-shf-gRNA-Xba1site-#1 | Generated by ligation into BbsI-digested pCFD5 vector of the annealed primers P17 and P18 |
| pCDF5-ptc-2xgRNA-CTerm: | Generated by Gibson Assembly into the BbsI-digested pCFD5 vector using the primers P19 and P20. |
| pCDF5-sd-gRNA1 | Generated by ligation into BbsI-digested pCFD5 vector of the annealed primers P63 and P64 |
| pCDF5- sd-gRNA2 | Generated by ligation into BbsI-digested pCFD5 vector of the annealed primers P65 and P66 |
| attb-U6-guide#1&2-nosCas9-attb (pHarvester) | Generated by 1) changing the direction of the SpeI insert in p*nos*-Cas9 by restriction and re-ligation. 2) Amplification of pCFD5gRNA1+2 with the Primers P21 and P22. 3) Restriction ligation of the fragment with ApaI-NheI into the vector generated in 1. |
| pSEED-sfGFP | Generated by 1) amplification of sfGFP fragments from pHD-sfGFP-ScarlessDsRed with primers P23+P24 and P25+P26 and 2) two-step restriction ligation of both fragments into pSEED-ALFAHA via its EcoRI+AvrII and Spe+SphI sites, respectively. |
| pSEED-ALFAHA | Generated by 1) Gibson Assembly of synthesized fragments “5' Alfa HA” and “3' Alfa HA” (from Genewiz) into HindIII-NdeI linearized pUC19 vector. 2) Amplification of the dsRED cassette from plasmid 80811 (Addgene) with primers P27 and P28 and restriction digestion cloning of the resulting fragment into the vector resulting from 1). |
| pSEED-mCherry | Generated by 1) amplification of mCherry fragments from pLOT-morphotrap with primers P29+P30 and P31+P32 and 2) subcloning into pSEED-ALFAHA via its EcoRI+AvrII and Spe+SphI sites, respectively. 3) Substitution of the 3xP3-dsRED cassette with the 3xP3-GFP cassette via restriction-ligation using HindIII and HpaI sites. |
| pSEED-EYFP:HA | Generated by 1) amplification of EYFP:HA fragments from a preexisting EYFP_HA plasmid with primers P55+P56 and P57+P58 and 2) two-step restriction ligation of both fragments into pSEED-ALFAHA via its EcoRI+AvrII and SpeI+SphI sites, respectively. |
| pSEED-OLLAS:HA | Generated by annealing of the primers P33+P34 and P35+P36 and ligation into pSEED-ALFAHA via its EcoRI+AvrII and SpeI+SphI sites, respectively |
| pSEED-Moontag:HA | Generated by triple ligation of: 1) PCR fragment amplified using P37 and P38 primers from a MoonTag plasmid and subsequently digested with EcoRI+EcoNI. 2) 3xP3-dsRed fragment obtained by digestion of pSEED-ALFA-HA with NheI and SpeI. 3) PCR fragment amplified using P39 and P40 primers from a MoonTag plasmid and subsequently digested with SpeI+EcoNI. The ligation product was then inserted into pSEED-ALFA:HA vector digested with EcoRI EcoNI. |
| pSEED | Generated by 1) digestion of pSEED-ALFAHA with NheI + SalI and 2) ligation into NheI + SalI-digested pBlueScriptIISK+ vector. |
| pSEED-T2A-Gal4 | Generated by 1) amplification of Gal4 fragments from pCMV-Gal4 (Dimitri Bieli) with primers P41+P42 and P43+P44 and 2) subcloning into pSEED-ALFAHA via its EcoRI+AvrII and SpeI+KpnI sites, respectively. |
| pSEED-ptc-ALFAHA | Generated by a 4-fragment assembly into NaeI+SfoI pSEED-ALFAHA digested vector. The fragments were 1) genomic Ptc fragment amplified with primers P45 and P46. 2) SEED-ALFAHA cassette obtained by Nae1+Sfo1 digestion from pSEED-ALFAHA. 3) genomic Ptc fragment amplified with primers P47 and P48. 4) genomic Ptc fragments amplified with primers P49 and P50. |
| pUC-GW-boi-mCherry-SEED | Generated by Gibson Assembly of the SEED-mCherry cassette amplified by primers P3 and P4 into the synthesized pUC19-GW-boi vector synthesized by Genewiz |
| pUC-GW-ihog-sfGFP-SEED | Generated by Gibson Assembly of the SEED-mCherry cassette amplified by primers P1 and P2 into the synthesized pUC19-GW-ihog vector synthesised by Genewiz |
| pUC-GW-shf-sfGFP-SEED | Generated by Gibson Assembly of the SEED-mCherry cassette amplified by primers P1 and P2 into the synthesized pUC19-GW-shf vector synthesized by Genewiz |
| pUC-GW-ihog-sfGFP-LOXP-SEED | Generated by amplification of the 3xP3-dsRED cassette of pSEED-sfGFP-LoxP with primers P1 and P2 and then subcloned into pUC19-GW-ihog via its NheI sites |
| pSEED-LoxPsfGFP | Generated by amplification of the 3xP3-dsRED cassette of pSEED-sfGFP with primers P51 and P52 and then subcloned into same plasmid via its NheI sites |
| pSEED-sd^GluGln^ | Generated by 1) amplification of sd genomic fragments using primers P67+P68 and P69+P70 and 2) two-step restriction ligation of both fragments into pSEED-ALFAHA via its EcoRI+AvrII and Spe+SphI sites, respectively. |
